# Compact RNA sensors for increasingly complex functions of multiple inputs

**DOI:** 10.1101/2024.01.04.572289

**Authors:** Christian Choe, Johan O. L. Andreasson, Feriel Melaine, Wipapat Kladwang, Michelle J. Wu, Fernando Portela, Roger Wellington-Oguri, John J. Nicol, Hannah K. Wayment-Steele, Michael Gotrik, Eterna Participants, Purvesh Khatri, William J. Greenleaf, Rhiju Das

**Author notes:** These authors contributed equally: Christian Choe, Johan O. L. Andreasson. Consortium author. All contributors are listed in Supplemental Table 1.

## Abstract

Designing single molecules that compute general functions of input molecular partners represents a major unsolved challenge in molecular design. Here, we demonstrate that high-throughput, iterative experimental testing of diverse RNA designs crowdsourced from Eterna yields sensors of increasingly complex functions of input oligonucleotide concentrations. After designing single-input RNA sensors with activation ratios beyond our detection limits, we created logic gates, including challenging XOR and XNOR gates, and sensors that respond to the ratio of two inputs. Finally, we describe the OpenTB challenge, which elicited 85-nucleotide sensors that compute a score for diagnosing active tuberculosis, based on the ratio of products of three gene segments. Building on OpenTB design strategies, we created an algorithm Nucleologic that produces similarly compact sensors for the three-gene score based on RNA and DNA. These results open new avenues for diverse applications of compact, single molecule sensors previously limited by design complexity.

## INTRODUCTION

Throughout biology, many macromolecular systems carry out complex calculations essential for life such as cell cycle regulation^1^, cell growth^2^, and tissue development^3^ but our ability to design comparably sophisticated biomolecular computers de novo remains primitive. Progress in the rational design of biomolecular computers would transform numerous fields – directed drug delivery, gene editors, and biosensors are technologies that all would benefit from computations at a microscopic scale in complex cellular environments. Sophisticated computations have been achieved in networks of interacting molecules,^4–12^ but such systems, which rely on a multitude of interacting parts at precise stoichiometries, are not always suited for complex *in vivo* environments – especially as the number of interacting parts grows to accommodate more complex computation. Computers that are instead based on single molecules might solve these issues and be capable of accurate complex computation in ambient cellular conditions or as ‘stand-alone’ computers outside cells.^13,14^ Furthermore, such single molecule computers might achieve thermodynamic efficiencies that outperform current electronic computers, potentially creating entirely new paradigms for low-energy computing technologies.^15,16^

Is there a limit to the functional forms that a single molecule can approximate? The behaviors of macromolecules have long been described through partition functions, which are ratios of two polynomials with non-negative coefficients. For example, expressions for hemoglobin behavior involve terms up to the fourth order in the partial pressure of oxygen and the concentration of protons and small molecules,^17,18^ and such expressions should be capable of near-arbitrary computations.^19^ Drawing motivation from this work and theorems derived for such positive rational polynomials^20,21^ (translation in **Supplemental Appendix 1**), a single macromolecule at equilibrium that is capable of binding input molecules and output molecules should be able to approximate any bounded polynomial (**Supplemental Appendix 2**). Here, we sought to more concretely explore the significance of these results by designing RNA-based approximators for functions involving polynomials with practical interest. The versatility of RNA as an allosteric biomolecular sensor is apparent in the diversity of natural “riboswitch” molecules that change their structure to alter downstream regulation upon binding an input molecule, which could be a drug, a metabolite produced by a downstream pathway, or a protein binding partner.^22^ Numerous examples of ‘tandem’ riboswitches exist that carry out computations involving multiple input ligands.^23^ RNA’s versatility is also demonstrated by the diverse synthetic elements, including designed allosteric riboswitches,^24,25^ therapeutics,^26,27^ and diagnostics^28^ that have been implemented with RNA as a substrate. Nevertheless, these prior efforts have been limited to ‘digital’ logic gates tested at extreme concentrations of two oligonucleotide inputs. To the best of our knowledge, molecular sensors designed to compute functions of multiple inputs across continuous ranges of concentrations – as is carried out by natural macromolecules – have not been successfully developed or tested.

As a driving application of a complex biomolecular calculation, we took inspiration from ratiometric gene signatures being discovered across diseases and host responses including sepsis, cancers, malaria, and pulmonary tuberculosis (TB).^29–32^ TB remains a major public health challenge worldwide, and the development of an accurate and accessible tool to discriminate active TB from latent TB and other diseases is a critical need. The World Health Organization has identified the need for a non-sputum-based triage test to identify individuals who require further testing.^33^ In this context, the use of a 3-gene transcriptional biomarker, including guanylate binding protein 5 (GBP5), dual specificity phosphatase 3 (DUSP3), and Krüppel-like transcription factor 2 (KLF2), has emerged as a promising signature for TB diagnosis.^34–36^ These genes collectively form a 3-gene signature referred as Sweeney3 or, in this manuscript, the ‘TB-score’. Sweeney et al. identified this combinatorial score based on blood messenger RNA (mRNA) expression levels, demonstrating its potential for discriminating active TB from other diseases. However, the complexity of the TB-score, which involves the quantity [GBP5][DUSP3]/[KLF2]^2^, currently requires expensive equipment involving quantitative RT-PCR, precluding routine usage at the point of care in resource-poor settings. A molecular sensor that could carry out the TB-score computation in samples after or during cell-free RNA amplification would enable diagnosis with potentially much lower cost and wider accessibility, but has not been achieved with nucleic acids, proteins, or cellular modalities.

To tackle this problem, we developed a set of crowdsourcing challenges for citizen scientists engaging in the Eterna videogame^37^. Prior work has demonstrated the ability of the Eterna community to solve RNA design tasks ranging from mRNA stabilization to design of small-molecule-activated RNA sensors achieving thermodynamic optimality.^38,39,39,40,13,14^ Here, we presented increasingly difficult challenges on the Eterna platform to build up to the final goal of designing a complex, multi-input sensor (**Figure 1**; **Supplemental Table 2)**. Within each challenge were a set of design puzzles, each representing a sub-problem of the overall challenge (**Figure 1b**). For example, in the pilot challenge of designing a single-input RNA sensor, one task was to design an ON-sensor while another task was to design an OFF-sensor. Both ON and OFF sensors accomplish the same goal of distinguishing the presence of an input RNA. The Eterna interface was extended to allow players to design an RNA for more than one condition simultaneously (**Figure 1c**) and to provide estimates of free energies of RNA folding in different conditions to give players rapid computational feedback.^41,42,43^ Although imperfect, these free energy estimates provide an approximation of the lowest energy secondary structure for a given sequence to guide player designs. After player designs were collected, they were synthesized and displayed on an Illumina sequencing chip for RNA-MaP (RNA on a massively parallel array)^38^ characterization. These experiments quantify the behavior of the player designs by measuring the affinity of the RNA sensor for a fluorescent output ligand across different input ligand conditions (**Figure 1d**-**f**). The results were then returned to the community, who, with this experimental feedback, were tasked with improving upon their previous results or tackling harder challenges (**Figure 1g**).

**Figure 1.**
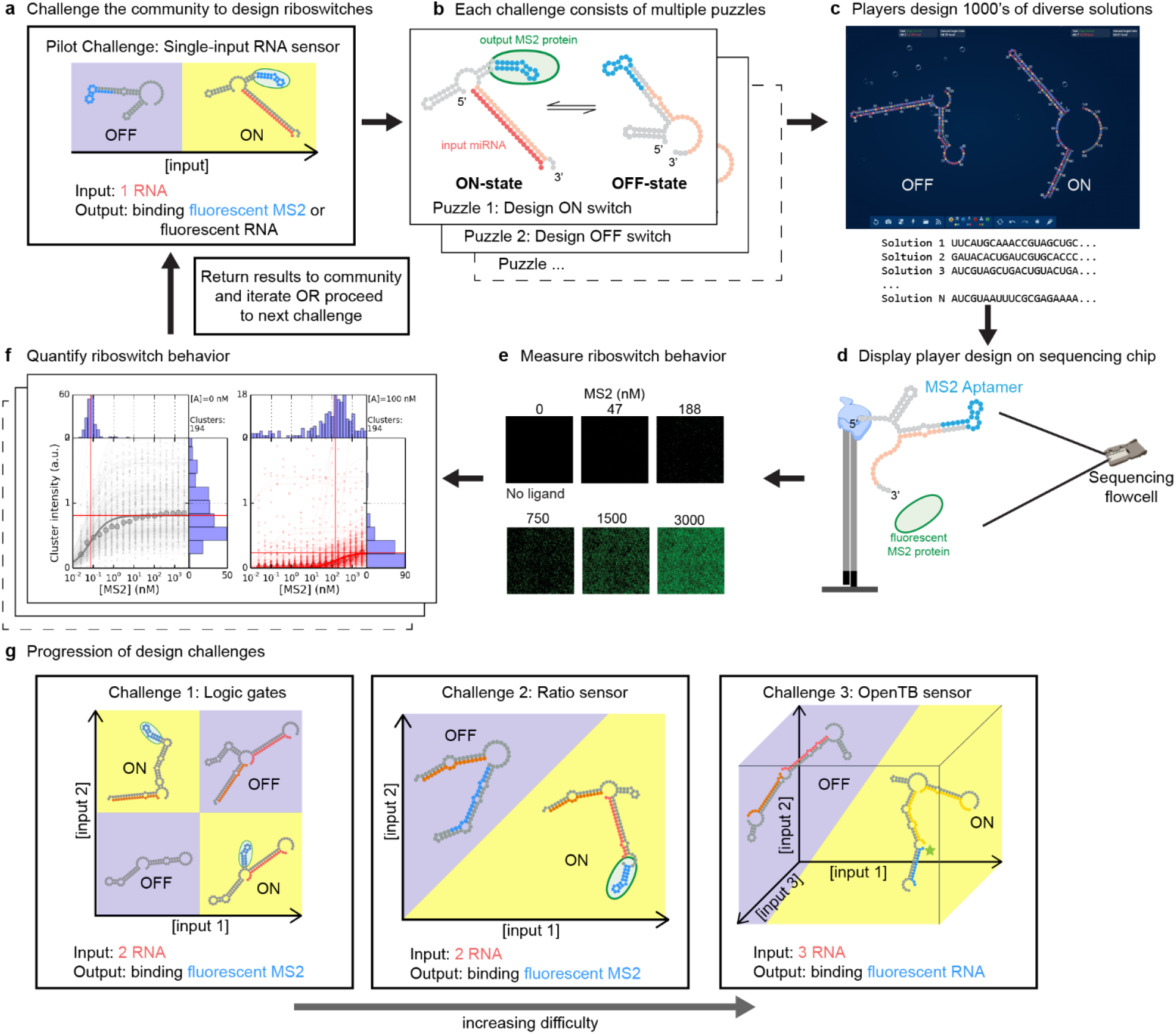
Pipeline for crowdsourced RNA sensor design and high-throughput testing. **a)** RNA sensor design challenges are presented to the Eterna community. [Input] represents the concentration of the input ligand. Concentration increases along the arrow direction. **b)** Each challenge consists of multiple puzzles such as designing an ON or OFF sensor for the specified inputs and outputs. **c)** Eterna interface enables players to design RNAs with two or more states. **d)** Player designs are synthesized by DNA array synthesis and converted to libraries ready for RNA-MaP characterization. **e)** Binding of fluorescent output reporter is quantified across all clusters at increasing reporter concentrations in the background of input molecules at fixed concentrations. Each cluster on the flow cell corresponds to a designed sequence. **f)** Binding data are quantified from multiple clusters for a single RNA sensor variant with the median fit shown. The data are then released to the Eterna community, and subsequent rounds of designs are solicited, or the next challenge is presented. **g)** The community is challenged with increasingly difficult design challenges gradually building up to the complex tuberculosis sensor. In **(a)** and **(g)**, yellow and blue coloring denote input conditions in which sensor response (binding of output ligand) is tighter or weaker than a specified threshold.

The Eterna design challenges culminated with the OpenTB challenge, which asked players to design an RNA sensor that could detect if the 3-gene TB-score is above the threshold that corresponds to active tuberculosis.^34^ This biomarker signal takes as input the mRNA concentrations of the genes GBP5, DUSP3, and KLF2. From each mRNA, we selected a short fragment to create simplified RNA inputs to the RNA sensor to be designed. Output fluorescence signal is generated by a fluorescently tagged RNA reporter engineered to bind to one of the possible states designed for the RNA sensor. Along with further testing with flow cytometry as an independent experimental readout, the results with OpenTB sensor design provide a proof of concept for using RNA sensors to detect a complex diagnostic signature.

Through these progressively more difficult challenges, Eterna players developed and documented new and productive RNA design strategies. We incorporated player-derived strategies into a Monte Carlo tree search algorithm, called “Nucleologic,” that allowed for the automation of the RNA sensor design process. Using Nucleologic, we generated candidate designs to compute this complex TB-score biomarker output signal. After experimentally testing a few selected designs, we identified a successful RNA and DNA sensor for the TB-score with performance comparable to top designs submitted by players. Nucleologic, which harnesses the human-inspired heuristics used by Eterna players, thus shows promise in expediting the process for designing nucleic acid sensors that can compute increasingly complex functions of multiple inputs.

## RESULTS

### Pilot challenge: Single-input RNA sensors

As an initial step towards more complex RNA design challenges, Eterna players were presented with a pilot challenge of designing single-input RNA sensors that responded to a separate RNA oligonucleotide. As a baseline, we sought to emulate or outperform prior work which has reported activation ratios as high as ∼900 for sensors of RNA oligonucleotides (albeit in cellular contexts).^44–46^ To connect this pilot round with the eventual challenge of designing TB-score sensors, we explored two signaling mechanisms (i.e., “outputs”). In the first mechanism, RNA sensors are turned ON by having binding of input RNA lead to display of an MS2 virus stem loop RNA structure, which recruits fluorescently-labeled MS2 virus coat protein (**Figure 2a**). This mechanism was chosen since Eterna players have previously designed RNA sensors with MS2 protein binding as an output signal.^38^ The input for these puzzles was a short RNA derived from hsa-miR-208a, a 22-nt miRNA (**Supplemental Table 3**) whose detection might aid in diagnosing cardiac hypertrophy^47^. Motivated by the final goal of developing sensors compatible with TB diagnosis, the second output mechanism involved hybridization of a fluorescent RNA reporter to a complementary sequence element in the sensor in the ON state. This output mechanism allows for incorporation in fluorescence-based or lateral flow-based diagnostic devices. Depending on the puzzle, players designed either an “ON” sensor where the RNA sensor fluoresces when bound to the input, or an “OFF” sensor where the RNA sensor fluoresces when not bound to the input.

**Figure 2.**
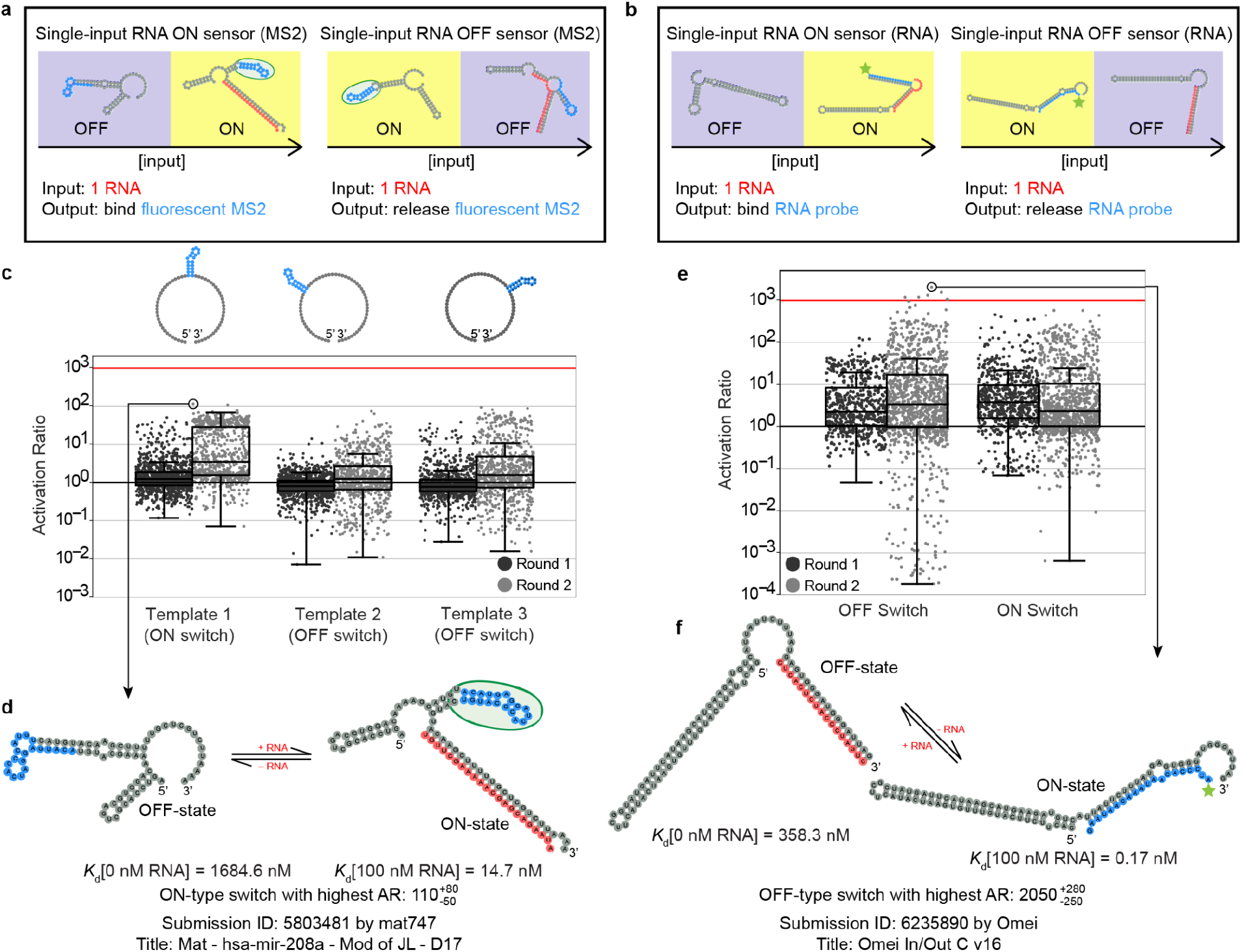
Pilot challenge: Single-input RNA sensors. The first challenge presented to the Eterna community was to design a single-input RNA sensor to detect the presence or absence of an input RNA oligonucleotide. **a)** One puzzle within the challenge involves designing the output to bind a MS2 coat protein fused to a fluorescent tag. **b)** The other puzzles involved binding or releasing an RNA oligonucleotide reporter conjugated to a fluorescent dye. **c)** Puzzle (a) results. (top) Players were constrained to three different templates for design with each template having a different MS2 hairpin location (top). Measured activation ratios across the architecture variants over two iterative rounds. **d)** Top player design of an RNA input/MS2 output ON-sensor for puzzle. **e)** Puzzle (b) results. Measured activation ratios for the ON and OFF sensors over two iterative rounds. **f)** Top player design of an RNA input/RNA output ON-sensor puzzle. In **(a)** and **(b)**, yellow and blue coloring denote subspaces in which sensor response (binding of output ligand) is tighter or weaker than a specified threshold. In **(c)** and **(e)**, the red horizontal line is the approximate maximum activation ratio that can be experimentally measured in RNA-MaP (1000). In **(d)** and **(f)**, RNA secondary structures were predicted using NUPACK.

Based on our prior work demonstrating the importance of widely exploring the relative placement of functional elements to achieve success,^14^ Eterna players were allowed to choose between three different templates that placed the MS2 aptamer sequence at different locations along the engineerable RNA molecule (**Figure 2c**). For the designs using an RNA reporter output, the flexibility of the NUPACK prediction algorithm allowed the position of the RNA reporter binding site to be left unconstrained. All designs were limited to 85 nucleotides in length.

For sensor designs that used MS2 binding as output signal, 3,369 and 2,319 player designs were characterized in Round 1 and 2 respectively, split across three different template options (**Figure 2c**). Round 1 presented players with only three templates while Round 2 introduced an additional template to increase the diversity of designs. The affinity of the RNA designs for their output molecules was then measured in the absence (0 nM) or presence (200 nM) of the input oligonucleotide. From the affinities for the output reporters, the activation ratio (AR) was calculated for each design by dividing the *K*_d_ [0 nM input] with the *K*_d_ [200 nM input]. The AR represents the fold change in the observed *K*_d_ of reporter binding between the OFF state (weak reporter affinity) and ON state (strong reporter affinity). Thus, larger AR values represent a sensor that better discriminates between the high and low input environments. We chose to use fold change in *K*_d_ to measure AR since it gives an unbiased and high signal-to-noise measure of performance by taking into account overall switch behavior across multiple output concentrations rather than at a single output concentration. In the limit that the output ligand concentration approaches zero, the fold change in observed *K*_d_ is equal to the AR values commonly reported in literature for switches (**Methods** and ref.^48^); because we can measure *K*_d_ values between 1 nM and 1 μM, AR values as high as 1000 can be precisely measured and are experimentally reproducible (**Extended Data Figure 1**). Player designs improved dramatically between the two rounds (**Figure 2c**). By refining previous submissions, players achieved designs with AR values close to or above 100 with the top design achieving an AR of 110 (error −50 values written in superscript and subscript correspond to one standard error, derived from fits to log *K*_d_, which give log_10_ AR of 2.06±0.22) (**Figure 2d**). Additional design refinement in a third round did not further improve AR values (**Extended Data Figure 2**).

Motivated by excellent performance in MS2-based output problems, 1,237 and 2,118 player designs were collected over two rounds for puzzles with RNA-based output more relevant for the TB-score sensors **(Figure 2e,f**). Round 1 used an 18-nt input and a shorter 10-nt reporter oligonucleotide, while Round 2 used a 17-nt input and a longer 20-nt reporter oligonucleotide. The reporter length was increased due to community feedback suggesting it was too difficult to design for a short output binding site (**Supplemental Table 3**). With a longer reporter oligonucleotide, players achieved AR above 1000 with a maximum observed AR of 2050^+280^_−250_ (log_10_ AR of 3.31±0.06) **(Figure 2e,f**) for OFF sensors. The ON sensors achieved slightly lower AR values, with a maximum observed AR of 570^+54^_−49_ (log_10_ AR of 2.76±0.04). Overall, throughout the pilot challenge, players achieved activation ratios at or beyond our experimental detection limits and previously reported RNA-triggered sensors.

### Challenge 1: Logic gates

We next challenged the Eterna community to generate designs of RNA Boolean logic gates (Figure 3a). This first full-scale challenge (Challenge 1) builds off of the pilot challenge by incorporating one additional input. While the goal was to gradually provide the Eterna design community experience in designing more complex multi-input RNA sensors, logic gates are, in their own right, useful tools in synthetic biology and nanotechnology and can, in principle, be chained together to execute complex computations.

**Figure 3.**
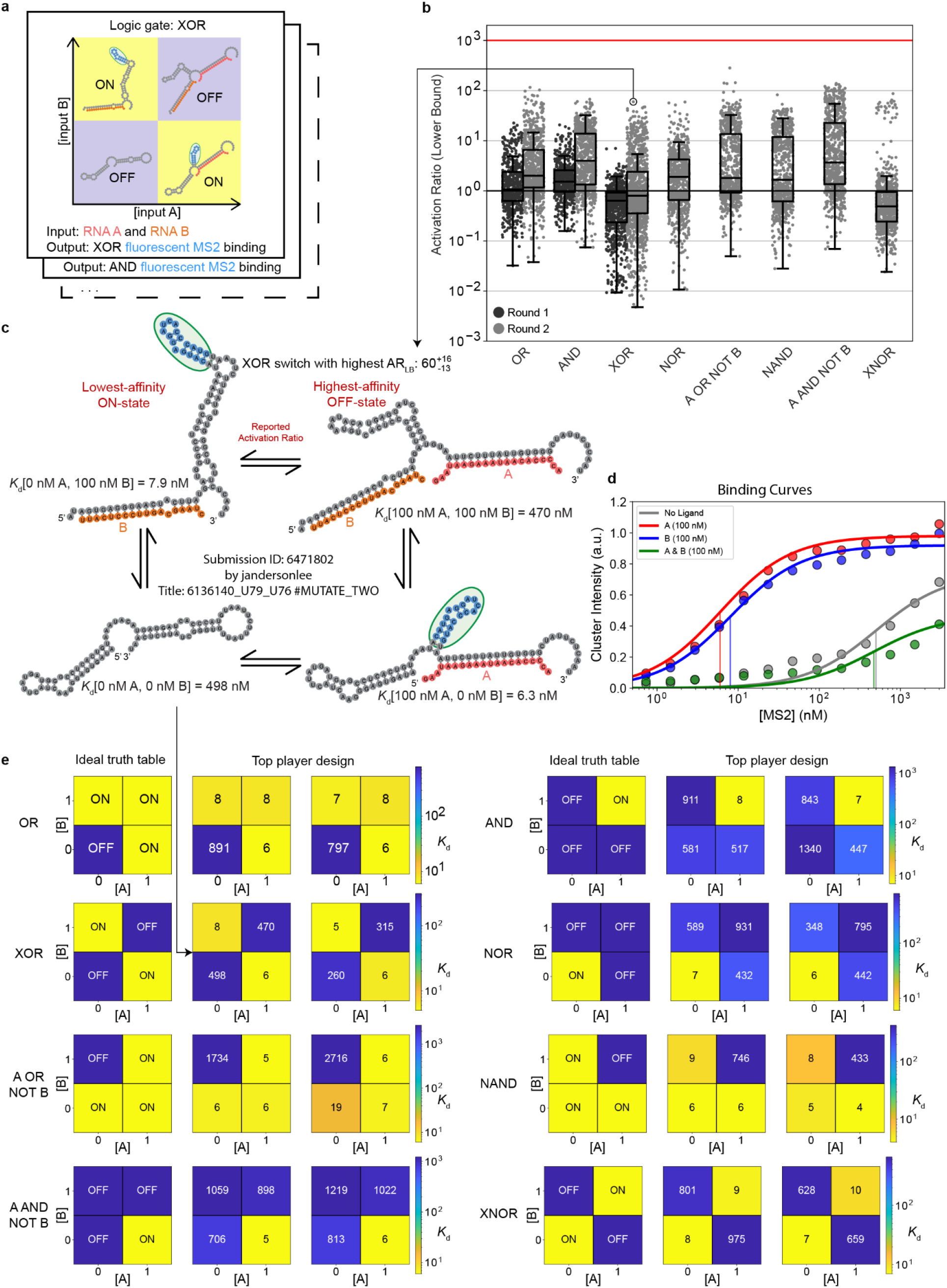
Challenge 1: Logic gates. Players were tasked with constructing all possible RNA sensors capable of computing Boolean logic gates of two different input RNAs denoted as A and B, with an output of binding a fluorescent MS2 coat protein. **a)** Each puzzle in this challenge corresponded to a different logic gate: AND, OR, NOR, XOR, NAND, XNOR, A OR NOT B, and A AND NOT B. **b)** Measured activation ratios (lower bound) of the eight different logic gates over two iterative rounds. The red horizontal line is the approximate maximum activation ratio that can be experimentally measured in RNA-MaP (1000). **c)** Top player design for XOR puzzle. RNA secondary structures were predicted using NUPACK. **d)** RNA-MaP binding affinity measurements of the player design in **(c)**. Vertical lines correspond to the *K*_d_ values. The points represent the median experimental fluorescence used to fit the binding curve. **e)** Ideal truth table and experimental results for each logic gate. The experimental data are from the top two designs from different Eterna player; *K*_d_ values given in nM. In **(a)** and ideal truth tables in **(e)**, yellow and blue coloring denote input conditions in which sensor displays or does not display the MS2 hairpin. In **(e)**, 0 and 1 ‘binary’ values correspond to 0 and 100 nM concentrations of A and B.

All logic gates were designed to bind fluorescently tagged MS2 protein as the output signal. In response to player feedback, an MS2 “stamp” tool was added to Eterna. This enabled players to easily place the MS2 hairpin RNA sequence anywhere they wanted within their design, giving players more flexibility in the design process compared to the Pilot Challenge. Each design was tested under four conditions corresponding to the four different binary inputs of the logic gate where A and B represent the first and second bit respectively. The binary input of 0 corresponds to 0 nM, while a binary input of 1 corresponds to 100 nM, and the A and B sequences re-used sequences of the input and output oligonucleotides used in the Pilot Challenge (**Supplemental Table 3**). To evaluate performance for each design, a conservative ‘activation ratio lower bound’ (AR_LB_) was computed by calculating the ratio of the *K*_d_ values for the poorest affinity OFF state with the tightest affinity ON state, where the OFF states and ON states are the conditions where an ideal logic gate would return a 0 or 1 respectively. Prior work on single-molecule logic gates have achieved AR_LB_ values as high as 21 for OR, AND, and NOR gates^9^ and above 100 when coupled to additional components like DNA polymerases^49^. AR_LB_ values for XOR and XNOR from RNA, protein, and DNA systems have remained below 10.^9,25,50–55^ We sought to determine if similar or better values might be achievable with single-molecule RNA sensors designed on Eterna.

In Round 1 of the logic gate challenge, Eterna players were tasked with designing OR, AND, and XOR gates. These design tasks were expanded in the second round to include NOR, A OR NOT B, NAND, A AND NOT B, and XNOR for a total of eight logic gates, which cover all possible truth tables (up to permutation of A and B; Figure 3b). During the first round, the best of 1,892 player designs achieved AR_LB_ values near 20, and in the second round, the best of 6,244 player designs achieved AR_LB_ greater than 100. Of all the logic gates, XOR gate was the most difficult for Eterna players to design, with the majority of Round 1 designs having an AR_LB_ value < 1. Nevertheless, after Round 2, the top player designs for XOR and XNOR achieved AR_LB_ of 60^+16^_−13_ (log_10_AR_LB_=1.78±0.10) and 87^+28^_−21_ (log_10_AR_LB_=1.94±0.12), respectively **(Figure 3c,d)**. Many successful XOR and XNOR gates in Round 2 (notably the distinct high AR_LB_ population of XNOR in **Figure 3b**) were designed by modifying sequences for Round 1 AND and OR gates that experimentally gave hints of XOR or XNOR activity (**Supplemental Table 4**). This result suggests that carrying out multiple simultaneous challenges on Eterna can lead to productive cross-fertilization of solutions. For all 8 logic gates, top player designs successfully approximate the logic gate outputs with the best designs achieving AR_LB_ greater than single molecule logic gates previously reported in the literature **(Figure 3e)**.

### Challenge 2: Ratio sensor

With the aim of building up to the final challenge of sensing the 3-input TB-score, which involves multiplication and division of the concentrations of input RNAs, we challenged the Eterna community to design an RNA sensor capable of computing the ratio of the concentrations of two input molecules. Specifically, players were tasked with designing an RNA sensor to detect if the ratio of two input RNAs A and B is greater than 1/4 **(Figure 4a)**. The key idea behind this challenge was driven by a mathematical form guaranteed by equilibrium thermodynamics: if a sensor can be designed to have two mutually exclusive states, one state binding A (but not B), and another state binding B but not A, the relative population of the states will be proportional to [A]/[B] (**Methods**). If the relative energetics of the two states can be set to achieve equal populations at [A]/[B] = ¼ and to favor the A-binding state under high [A]/[B] conditions and the B-binding state at low [A]/[B] conditions, the sensor responds to the desired ratio.

**Figure 4.**
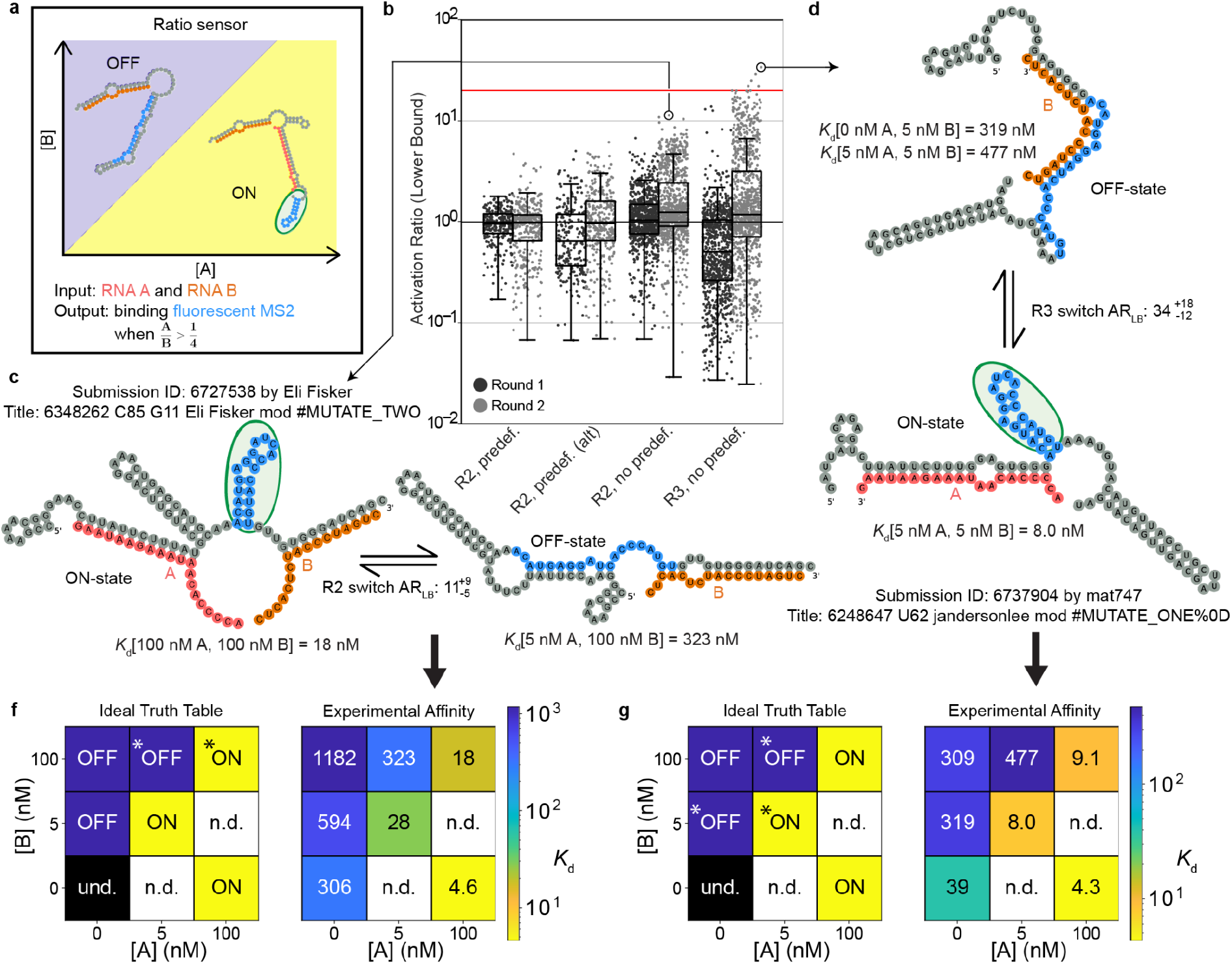
Challenge 2: Ratio sensor. **a)** Players were tasked with designing an RNA sensor that turned on when [A]/[B] > ¼, with binding of fluorescent MS2 coat protein as output signal. **b)** Measured activation ratios (lower bound) across different puzzles and rounds. The red horizontal line is the theoretical maximum for AR_LB_ of 20 for the tested conditions. **c)** Top player design from the R2 puzzles. **d)** Top player design for the R3 puzzle. **f)** Ideal truth table and the player design truth table for **(d)**. **g)** Ideal truth table and the player design truth table for **(c)**. The * symbol in the ideal truth table of **(f)** and **(g)** indicates the *in silico* conditions simulated in Eterna puzzles. Undefined = und., not defined = n.d. RNA secondary structures in **(c)** and **(d)** were predicted using NUPACK. In **(a)** and ideal truth tables in **(f-g)**, yellow and blue coloring denote input conditions in which sensor displays or does not display the MS2 hairpin.

Several different puzzles were created to explore which in silico constraints might yield the most performant ratio sensors (**Figure 4b**). The “R2” puzzles tasked players to design a sensor that exhibited the desired behavior at two different simulated input concentrations, while an “R3” puzzle tasked players to design a sensor that exhibited the desired behavior at three different input concentration combinations. The simulated design conditions for the R2 puzzle were 5 nM A & 100 nM B and 100 nM A & B, corresponding to A/B of 1/20 and 1. Design conditions for the R3 puzzle added a third condition, 0 nM A & 5 nM B, corresponding to [A]/[B] of 0. The hypothesis for R3 was that by having an additional OFF condition constraining the puzzles, designs submitted would be more robust across all A and B concentrations tested experimentally. Furthermore, we sought to understand whether allowing players to each explore a wide set of binding sites for A and B might be better than focusing the Eterna community’s attention on specific sets of predefined binding sites. We deployed three sets of R2 puzzles to test this idea (“predef.”, “predef. alt”, and “no predef.”, **Figure 4b**). Finally, to test generality of player design strategies, we used different A and B sequences here than in Challenge 1, using input and output sequences from different rounds of the Pilot Challenge (**Supplemental Table 3**).

The designs were experimentally tested across a total of seven conditions, expanding the two or three conditions presented to Eterna players (**Figure 4f,g**), over two rounds with 2,254 and 2,534 designs tested, respectively. AR_LB_ was again computed as a worst-case ratio of the highest-affinity OFF state with the lowest-affinity ON state with respect to all seven test conditions; the best possible theoretical AR_LB_ value achievable was 20. Across the two design rounds, many player designs achieved AR_LB_ values greater than 10. The top player design from R2 puzzles gave an AR_LB_ of 11^+9^_−5_ (log_10_AR_LB_ 1.06 ± 0.26) (**Figure 4c**). These top designs came from the R2 puzzle without predefined binding sites, which overall led to significantly better AR_LB_ than the two puzzles that constrained designs with predefined A and B binding sites. This result supported the principle that wide varieties of design patterns should be explored during the design process, which was also supported by our prior work on small molecule sensors^14^ as well as the previous challenges in this study (**Figures 2**-**3**). While the average performance in R3 is slightly worse than in the R2 puzzles, the best overall submissions were from the R3 puzzle, supporting our hypothesis that an additional simulated design condition would favor better solutions (**Figure 4b**). The top R3 designs achieved experimental truth tables similar to the ideal truth table across all conditions and a top AR_LB_ of 34^+18^_−12_ (log_10_AR_LB_ = 1.53 ± 0.18) (**Figure 4f-g**). Interestingly, many designs from this round came from strategies developed by Eterna players in the previous challenges and developed further in the following challenge (**Supplemental Table 4**).

### Challenge 3: OpenTB sensors

After successfully creating two input sensors for logic gates and a ratio function, we challenged Eterna players to design an RNA sensor to compute the 3-gene TB-score for active tuberculosis. In clinical studies, active tuberculosis correlates with expression of three gene mRNAs (GBP5, DUSP3, and KLF2; here called A, B, and C) in which [A][B]/[C]^2^ is greater than or equal to 1/16 (**Methods**). In the OpenTB challenge, we envisioned that both positive and negative sensors for this signature would be clinically useful either separately or in combination for a more robust diagnostic. We therefore challenged the Eterna community to design RNA sensors to address the full TB-score calculation: [A][B]/[C]² > 1/16 (“INC” to detect when the TB-score is above the threshold) or [A][B]/[C]² < 1/16 (“DEC” to detect when the TB-score is below the threshold) (**Figure 5a**). Additional puzzles were presented to let players experiment with designing simpler intermediate RNA sensors related to this final sensor **(Supplemental Table 2; Extended Data Figure 3**).

**Figure 5.**
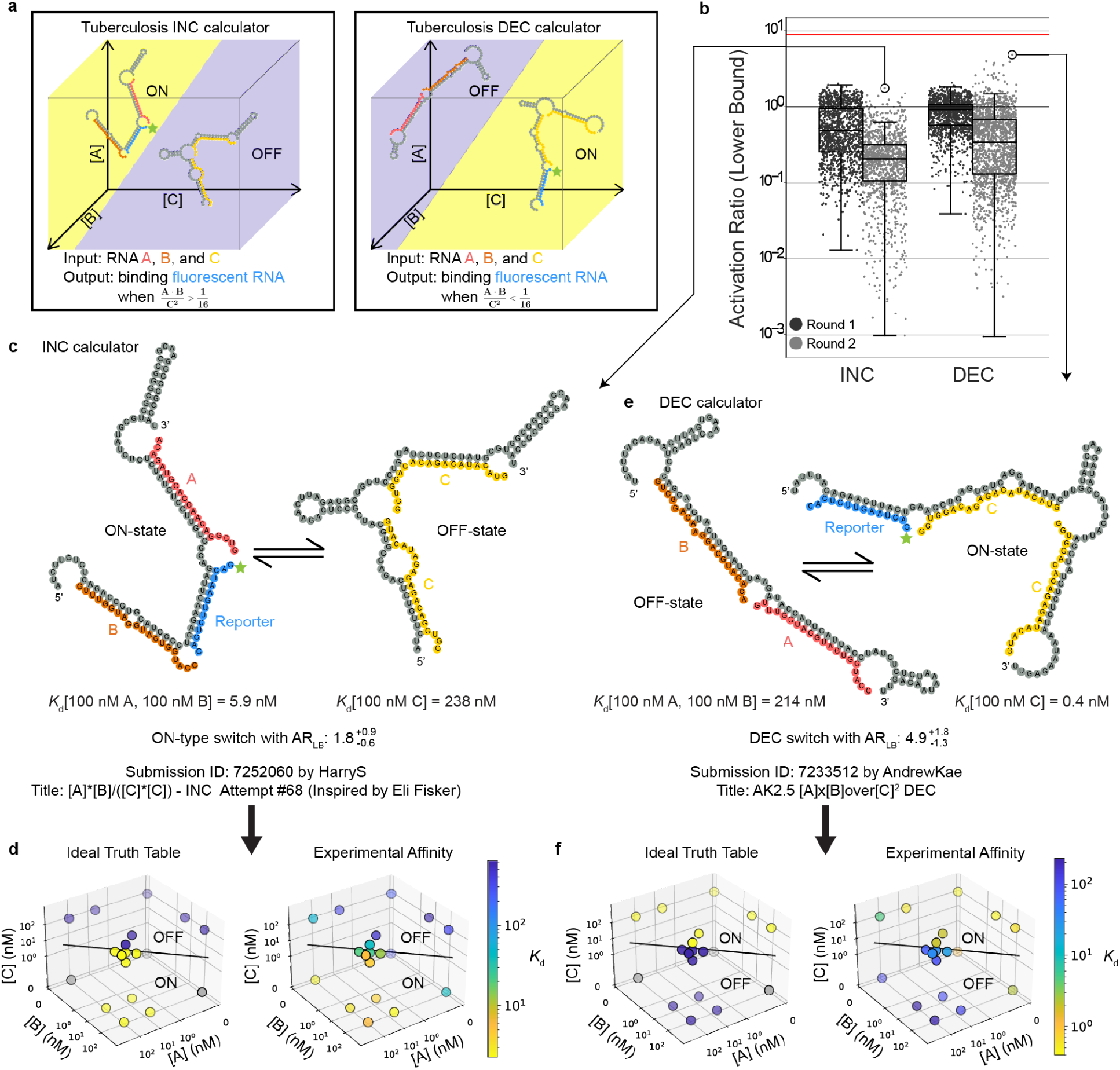
Challenge 3: OpenTB sensors. The three inputs are designated as A, B, and C with the output being a fluorescent RNA reporter. **a)** Players were tasked with designing an “INC” TB-score sensor with the fluorescent RNA reporter binding when [A][B]/[C]^2^ > 1/16 (left), or a “DEC” sensor binding fluorescent RNA reporter when [A][B]/[C]^2^ < 1/16 (right). **b)** Measured activation ratios (lower bound) for INC and DEC designs across two iterative rounds. The red horizontal line is the theoretical maximum for AR_LB_ of 9. **c)** Best INC player design from Round 2. **d)** Ideal truth table and experimental truth table for INC design in **(c)**. **e)** Best DEC design “AK2.5” from Round 2. **f)** Ideal truth table and experimental truth table for DEC design in **(d)**. RNA secondary structures in **(c)** and **(e)** were predicted using NUPACK. In **(d)** and **(f)**, the separatrix plane for [A][B]/[C]^2^ = 1/16 appears as a line due to the chosen view of the 3D plot, and gray points in ideal truth tables are undefined with respect to the three-gene TB-score.

Players were asked with designing RNA sensors that exhibit two mutually exclusive states, one state binding a single copy of A and B but no C strands, and the other state binding two copies of C but no A or B. The puzzles involved four simulated design conditions: 100 nM A & 100 nM C, 100 nM B & 100 nM C, 50 nM A&B & 300 nM C, 50 nM A&B & 100 nM C. For all conditions, 25 nM of the RNA output reporter R (initially chosen to be the same as prior Challenge 2) is present. These four conditions correspond to values of the TB-score ratio equal to 0, 0, 1/36 and 1/4 respectively. The first two conditions that keep A or B as 0 nM ensure player designs can properly bind A and B and will treat them as functionally interchangeable since they are both multiplied on the numerator. This helps to avoid designing an [A]/[C] or [B]/[C] sensor. Having the sensor switch its favored state between the last two conditions ensures that binding of C competes with the binding of A and B; requiring two copies of C bind cooperatively ensures the sensor’s output involves [C]^2^ in the denominator. Altogether, these four simulated solution conditions establish strong boundary conditions for player designs. The standard formula for chemical equilibrium then requires the ratio of the sensor’s two states to be proportional to [A][B]/[C]^2^ and helps ensure that the sensor should sustain its desired behavior at other concentrations of A, B, and C (**Methods**), an assumption we also tested experimentally below. Throughout the OpenTB challenge, players also had access to simulation plots which provided *in silico* NUPACK predictions at a whole suite of input concentrations ranging from fM to mM to help players refine sequences beyond the four conditions presented in Eterna **(Extended Data Figure 4)**. In addition, after each round of experiments, the Eterna community was given a detailed PDF summary of each design showing the binding curves of the reporter RNA under all conditions tested in RNA-MaP (example in **Extended Data Figure 7**).

For Round 2, the 20-nt input sequences were changed after players selected new fragments from the TB-score genes GBP5, DUSP3, and KLF2 based on BLAST analysis **(Supplemental Table 3)**. Also, based on player recommendation, the RNA output reporter was lengthened from 10-nt to 14-nt to create more binding potential. Finally, a “Freeze Mode” was added to Eterna at player request which allows players to modify their RNA sequence without triggering NUPACK computation, which required over a minute on most player computers.

A total of 2,694 Round 1 designs were tested at 13 different combinations of A, B, and C input concentrations, while 2,818 Round 2 designs were tested against 19 different combinations to more fully explore the phase space of behaviors **(Supplemental Table 5)**. Under each of these conditions, full binding curves were derived measuring the effective *K*_d_ of the output signaling reporter to the RNA molecule. For each design, AR_LB_ was computed as a ‘worst-case’ metric, similarly to the previous challenges, as the ratio of the highest-affinity OFF state with the lowest-affinity ON state with respect to all experimentally tested conditions; a perfect sensor would achieve AR_LB_ of (1/4)/(1/36) = 9. The best INC design achieved AR_LB_ of 1.8^+0.9^_−0.6_ (log_10_ AR_LB_ = 0.25±0.18; **Figure 5c**). Worse INC designs were observed in Round 2 compared to Round 1. This decrease in AR_LB_ may be due to the increased number of conditions tested, which increases the likelihood of observing errors in the sensor that are sensitively captured by AR_LB_ (**Figure 5b**). The DEC design challenges were more successful, with player designs like AK2.5 reaching an AR_LB_ of 4.9^+1.8^_−1.3_ (log AR_LB_ = 0.69±0.13) (Fig 5b**,d**). These values are affected by experimental uncertainties in some of the 19 test conditions, likely leading to artificial suppression of AR_LB_. When focusing specifically on the four A, B, and C conditions that were simulated in the Eterna puzzle, this same AK2.5 sensor gives an AR_LB*_ of 11.5^+33^_−2.5_ (log_10_ AR_LB*_ = 1.06±0.11; see **Extended Data Figure 5**), agreeing with the maximum value of 9. To test if the improvements might be due to the updated input and output RNA sequences, Round 3 repeated the challenges but reverted these sequences to the original Round 1 sequences (**Supplemental Table 2**); indeed, this round led to sensor performance as poor as Round 1 (**Extended Data Figure 6**). For further independent evaluation across a broader range of input conditions, the top INC and DEC designs were carried forward to flow cytometry measurements, described next.

### Flow Cytometer Characterization of the Best OpenTB sensor

As an independent and more thorough test of functional accuracy, the top-scoring player designs from the OpenTB challenge were selected for characterization across a wide range of input conditions using flow cytometry (**Figure 6a**). This orthogonal measurement of fluorescence response was achieved by first attaching the sensors to the surface of magnetic beads and then incubating with a fluorescent RNA reporter (30 nM) at different input concentrations of A, B and C input RNA molecules (**Figure 6b**). Because some of the [A][B]/[C]^2^ conditions in the flow cytometry experiments approach closely to the separatrix 1/16, the best achievable AR_LB_ metric would approach 1 even for perfect designs and not be useful for ranking. We therefore ranked designs by a metric more common in diagnostic characterization, the area under the receiver operating characteristic curve (AUROC), which varies the output fluorescence threshold and computes specificity and sensitivity. In agreement with RNA-MaP measurements, DEC sensors outperformed INC sensors in flow cytometry. A particularly notable DEC sensor with excellent performance at low concentrations of A, B, and C input RNA molecules was a close homolog of AK2.5 named AK2.2 (**Figure 6c** and **Extended Data Figure 9**), which achieved AUROC of 0.935 under these test conditions (**Extended Data Figure 8**). AK2.2 was carried forward for more detailed testing across 144 different conditions (**Figure 6c**). Overall, AK2.2 achieved AUROC 0.959 across these conditions (95% confidence interval CI, 0.930–0.988; **Figure 6d**). Across the entire input-space volume that was experimentally tested, AK2.2 was able to properly categorize points as positive or negative with a specificity of 89.6% and a sensitivity of 89.5% at a threshold chosen to maximize the sum of the specificity and sensitivity (red line, **Figure 6e**). The sensor performance was expected to be best at the highest input oligonucleotide concentrations, where the assumption that either A and B or two copies of C bound would best hold, without states with fewer input oligos bound. Indeed, at concentrations of [A]+[B]+[C] > 170 nM, the sensor performance is visually clearer (black points in **Figure 6d**) and AUROC increases to 0.979 (95% CI, 0.948–1.0) for A, B, and C in this high input concentration range.

**Figure 6.**
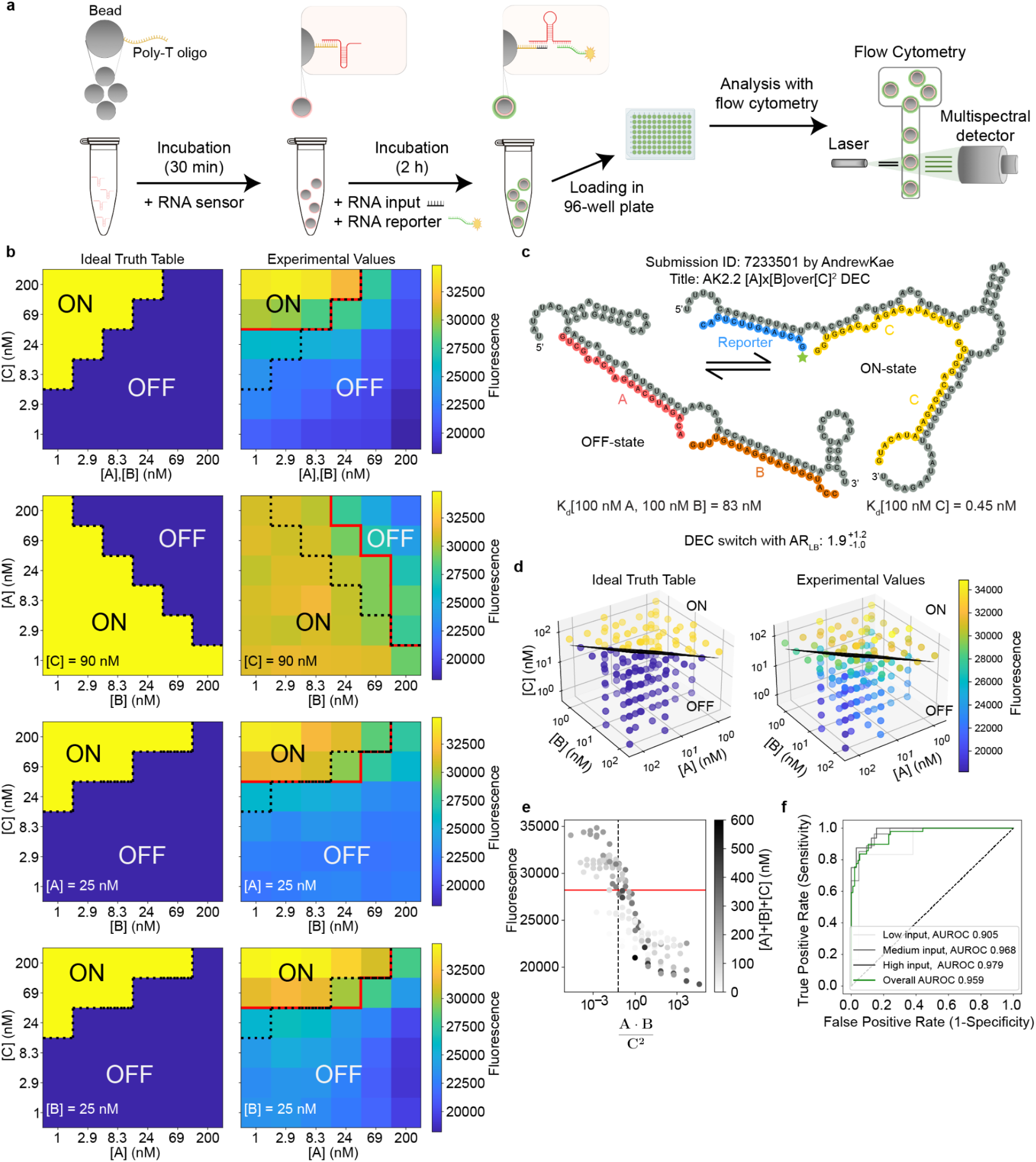
Flow cytometer evaluation of the top performing DEC tuberculosis sensor. **a)** Experimental pipeline for testing an RNA sensor with flow cytometry. 1 micron diameter beads coated in (dT)_25_ were incubated with the RNA sensor and then washed to remove unbound RNA. The beads were then incubated at various conditions and fluorescence measured in the flow cytometer. **b)** Ideal and experimental truth tables across various conditions: (top to bottom) A,B against C with A equal to B; A against C with B = 25 nM; A against B with C = 25 nM; and B against C with A = 25 nM. The red line corresponds to the optimal fluorescent threshold to approximate the intended logic function to diagnose active tuberculosis. The dotted black line corresponds to the ideal threshold for diagnosis. **c)** Best sensor AK2.2 tested by flow cytometry; RNA secondary structures predicted using NUPACK. **d)** 3D scatter plot of the ideal truth table (left) and experimental values (right) with the dividing plane at [A][B]/[C]^2^ = 1/16. **e)** Distribution of measured fluorescence of design AK2.2 across different values of [A][B]/[C]^2^. The black dashed line represents the tuberculosis diagnostic threshold at [A][B]/[C]^2^ = 1/16 and the red line represents the best fluorescence cutoff to diagnose active tuberculosis. **f)** Receiver operating characteristic (ROC) curve of design AK2.2. ‘Low input’, ‘medium input’, and ‘high input’ correspond to computing AUROC for data points where [A]+[B]+[C] values are <75 nM, 75-170 nM, and >170 nM, respectively. In **(d)**, the separatrix plane for [A][B]/[C]^2^ = 1/16 appears as a line due to the chosen view of the 3D plot.

### Computational design of RNA and DNA sensors for the TB-score

Starting with early design rounds on two-input Boolean logic gates, Eterna players derived a sequence-independent heuristic for sensor design named Domain Matching Secondary Structure Design (DMSSD; see **Supplemental Table 4**). DMSSD deploys a constraint-driven method using predefined domains to design secondary structures. The method “chunks” the RNA sequence into domains where each domain is associated with another complementary or near-complementary domain in the sequence to facilitate intramolecular interaction via secondary structure. This method is itself an improvement upon a common player strategy called kernel attractors (**Supplemental Table 4**) where a domain in the designed sequence will be “attracted” to the output and input oligomers due to complementary sequences. While the kernel attractor strategy requires fine tuning the interaction strength of one domain with two or more sequences, DMSSD simplifies the process by focusing on designing interactions between a domain and its complementary domain and relying on mutual exclusion of interleaved stems in RNA secondary structure.

**Figure 7a** shows an example of employing DMSSD to design an RNA sensor to detect two input RNAs, A and B, along with an output RNA R. First the RNA sensor sequence is partitioned into several domains: A’, A”, B’, B”, R’, and R”. Domains A’ and B’ were designated to harbor complementary sequences to input A and B respectively. For the output, domain R is complementary to a fluorescently tagged RNA reporter. In addition to these domains, extra domains were added that are complementary to an existing domain such as A” which is complementary to A’. This allows for complex secondary structure rearrangement in the absence and presence of the input ligands due to the network of complementary regions in the RNA by interweaving the domain locations. In particular, the R’-R’’ stem cannot occur simultaneously with the A’-A’’ or B’-B’’ pairing. By altering the complementarity between domains and the order of domains, it is possible to alter the RNA sensor’s response to fit the boundary conditions of a user-defined function f([A],[B]).

Inspired by these design principles from Eterna players, we created Nucleologic, a Monte Carlo tree search algorithm for automating the design of complex RNA sensors (**Figure 7b-c**). The input to Nucleologic is the set of domains that make up the single strand nucleic acid sequence as well as the order of the domains. Typically, a domain with a sequence complementary to the input and output RNAs is included with extra domains that are filled with N’s. During sequence optimization, Nucleologic can perform two types of moves to the sequence: domain mutation, which mutates the sequence within the domain, or domain move, which moves the location of a domain. The choices of which intermediate solutions to carry forward are made based on Monte Carlo tree search, a classic automated game playing strategy.^56,57,58^ By posing the problem as a game with DMSSD-inspired moves, Nucleologic optimizes the activation ratio and fold-change between the ON-states and OFF-states defined by the user calculated using NUPACK. Using Nucleologic, we computationally designed several hundred DEC and INC RNA sensors for the TB-score, screened four designs by flow cytometry, and carried forward one of these HP_MCTS_130 for testing across 144 input conditions (**Extended Data Fig. 10** and **Figure 7d**). While HP_MCTS_130 does not perform as well as AK2.2, it has a lower baseline fluorescence and can accurately discriminate between the ON and OFF states when [C] is low (**Figure 7e-f**). Across the 144 points in the input-space volume that were experimentally tested, HP_MCTS_130 was able to properly categorize points as ON or OFF with a specificity of 78.5% and a sensitivity of 69.5%, and AUROC of 0.900 (95% CI 0.852–0.947). In contrast to the Eterna design AK2.2, this Nucleologic RNA sensor achieves better performance at lower input concentrations: at [A]+[B]+[C] of 70 nM or lower, HP_MCTS_130 gives an improved AUROC of 0.932 (95% CI 0.839–1.0). To test the generality of Nucleologic, we then tested two sensors for computing the TB-score based on DNA instead of RNA (**Extended Data Fig. 10**). Experimental results from flow cytometry demonstrated that one of these, named MCTS_DNA_DEC, switches appropriately when A and B are varied against C with a specificity of 78.47% and sensitivity of 69.47% (**Figure 7f,g**). The DNA sensor achieves an AUROC of 0.899 (95% CI 0.852–0.947) (**Figure 7f**).

**Figure 7.**
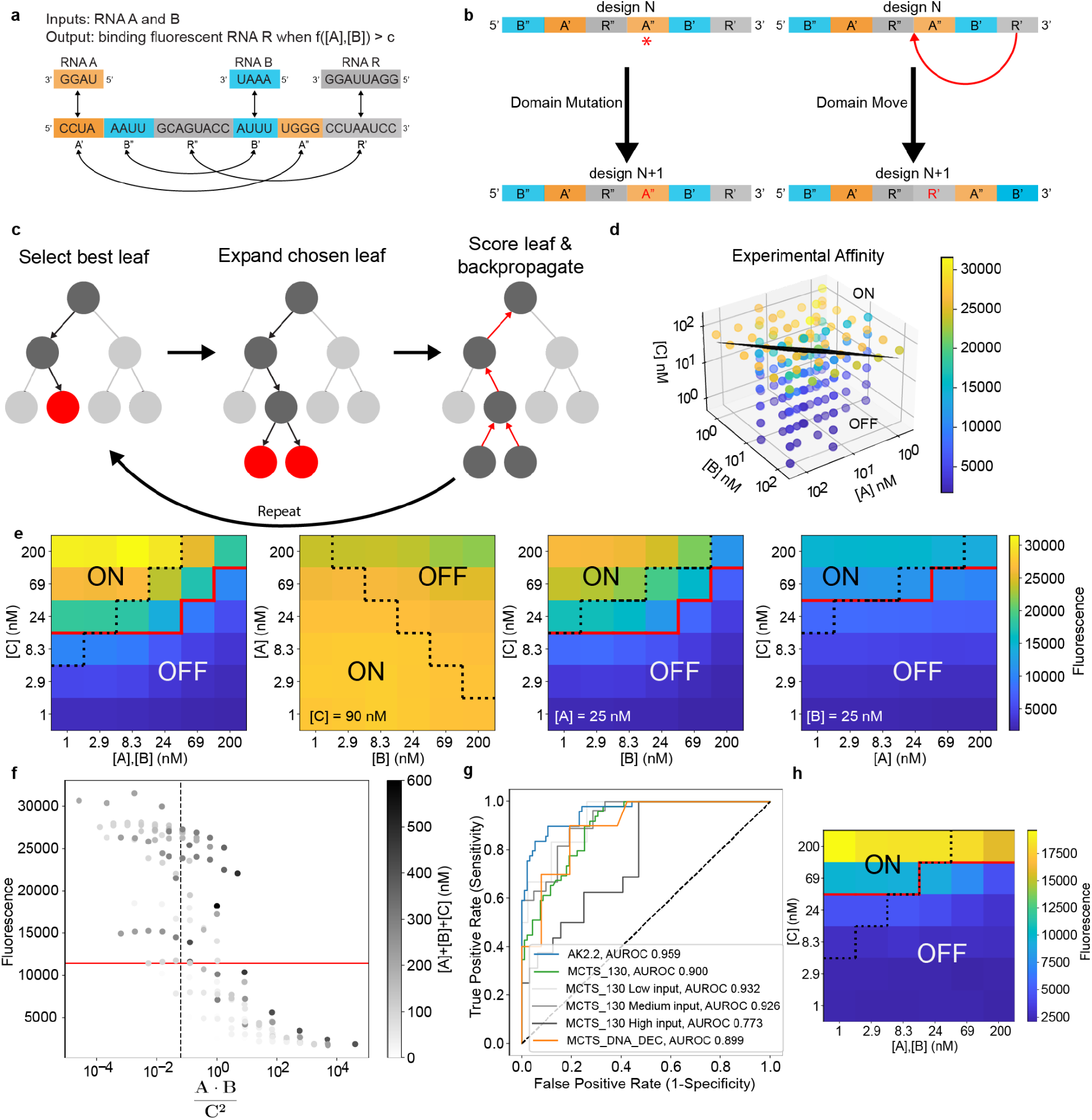
Nucleologic automates the design of complex nucleic acid sensors by integrating player inspired heuristics with a classic game play strategy, Monte Carlo tree search. **a)** Schematic example demonstrating the use of Domain Matching Secondary Structure Design (DMSSD) for a toy problem involving two input RNAs designated as A and B and one output RNA reporter designated as R. The design is grouped into blocks of domains each with a label that designates to which domain it is complementary. For example, A’ is complementary to the input RNA A as well as to A”. The domains are interwoven such that the domains compete with each other for the formation of base paired stems. **b)** Nucleologic performs two different types of modifications to sensor sequences: 1) “Domain mutation” creates a random point mutation, and 2) “Domain move” removes the domain and reinserts it into a new location. **c)** The Nucleologic algorithm consists of three steps: selecting the best leaf node based on the Upper Confidence bound for Trees (UCT) score, expanding the leaf node by generating children using the modifications in **(b)**, and scoring the leaf and backpropagating the value, updating the UCT score of all nodes in the path back to the root node. **d)** Example of a design of a DEC sensor HP_MCTS_130, designed using Nucleologic. **e)** Experimental truth tables measured at each condition in **(d)**. **f)** 3D scatter plot of the ideal truth table (left) and experimental values (right) with the dividing plane at [A][B]/[C]^2^ = 1/16. **g)** Receiver operating characteristic (ROC) curve of the Nucleologic RNA sensor HP_MCTS_130 over all tested conditions (green). Bins 1 to 3 correspond to computing the AUROC for data points where [A]+[B]+[C] are <75 nM, 75-170 nM, and >170 nM. ROC curves for AK2.2 (blue) and the Nucleologic DNA sensor MCTS_DNA_DEC (orange) are shown for comparison. **h)** DNA DEC tuberculosis sensor designed using Nucleologic. Experimental truth table for A,B (at equal concentrations) against C. In **(e)** and **(h)**, the red line corresponds to the optimal fluorescent threshold to diagnose active tuberculosis. The dotted black line corresponds to the ideal threshold for diagnosis.

## DISCUSSION

RNA and DNA are ideal substrates for designing function approximators due to the ease of large-scale nucleic acid synthesis, availability of computational modeling methods for predicting nucleic acid structure, and increasing throughput of experimental evaluation methods. Nevertheless, the complexity of functions achieved by single molecule nucleic acid sensors has been limited. Here, starting from simple single-input RNA sensors, a community of Eterna citizen scientists successfully designed more complex multi-input RNA sensors including all possible logic gate sensors, two-input ratio sensors, and, in the OpenTB challenge, sensors of the 3-gene TB-score [GBP5][DUSP3]/[KLF2]^2^ for diagnosing active tuberculosis. In all challenges, the activity ratios of Eterna sensors approached the limits of our experimental assays or the theoretical upper bound for the sensors, and exceeded activation ratios achieved by prior studies. The performance of Eterna sensors was particularly striking given their compact lengths – the 85 nucleotides of the TB-score sensors presented binding sites for the 14-nt reporter as well as GBP5, DUSP3, and two KLF2 segments (20-nt each). In addition, the sensors were robust to transferred from repurposed Illumina sequencers (for RNA-MaP) to display on beads for more complete characterization by flow cytometry. Driving the success of these compact, near-optimal designs were novel Eterna strategies such as the kernel attractor and DMSSD heuristics. By incorporating these strategies into a Monte Carlo tree search, we created Nucleologic to automate the design of complex nucleic acid sensors. Our work demonstrates the continuing utility of citizen science-based crowdsourcing integrated with iterative, high-throughput experimental evaluation to design complex RNA molecules.

These results suggest a route to developing nucleic acid sensors for multi-gene signatures for diseases beyond active tuberculosis, including septic shock, cardiovascular disease, vaccine-induced immunity to malaria, and cancers.^30–32,3630–32,36,4730–32,36^ However, there are limitations to the current proof of concept. Eterna and Nucleologic currently cannot handle sensor designs with more than 6 inputs due to the impractical factorial scaling of computational time with the number of interacting strands.^43^ This limitation currently precludes automated design of, e.g., the 10-gene signature for sepsis.^30^ Furthermore, in future efforts, it will likely remain necessary to experimentally screen multiple designs; current automated designs have generally acceptable performance (AUROC of 0.9) but are worse at extreme input gene concentrations, presumably due to inaccuracies in available modeling packages. Importantly, the application of single molecule sensors for low-cost diagnostics will require enzymatic amplification of gene segments and the readout of reporter binding in inexpensive platforms. Such technologies have been developed for single-gene sensors^59–61^ but remain to be demonstrated for the more complex multi-gene sensors like the ones described here.

There are more general limitations to this work. Most fundamentally, we only designed single-molecule sensors for up to three distinct inputs and for systems restricted to two states. The space of functions able to be approximated by a molecular sensor is much larger, comprising the space of all positive rational polynomials. This space theoretically allows for approximation of any continuous function in the nonnegative quadrant with nonzero leading homogeneous term (**Supplemental Appendices**). Designing systems that have more than two states and binding of more than three inputs would allow for sensing of more complex functions, with more elaborate contours like ellipses or the piecewise linear functions encoded by artificial neural networks,^19^ but has not been carried out. Last, for applications involving low-energy computing or embedding computation in nucleic acid therapeutics, thermodynamic reversibility of sensors would enable repeated and continuous use in real world settings. Such reversibility appears feasible through the approaches described here and has been demonstrated for RNA sensors of single small molecules^14^ but has not been demonstrated for multi-input sensors.

## MATERIAL AND METHODS

### Eterna online interface

The design of RNA molecules in Eterna has been described previously.^14,37^ In this work, the interface was further improved to allow for visualization, calculations, and design using several strands of RNA. Multi-stranded folding calculations from the NUPACK folding package^43^ were integrated into Eterna, thereby providing players with computational feedback during the design process. For designs utilizing RNA reporter output, the RNA reporter binding site is fully unconstrained in terms of binding site location, and the simulated reporter concentration was set to 100 nM (single input sensors) or 25 nM (OpenTB). The puzzle interface provides visualization of the secondary structure of the complex with highest probability under each simulated condition. All RNAs were constrained to have no repeat of any four nucleotides and uniform lengths of 85 nucleotides to aid synthesis. Only the RNA in/MS2 out designs are 77 nucleotides. Wet-lab experimental scores were converted to numbers between 0 and 100 and were based on either activation ratios (AR) or, for multiple state problems, lower bound activation ratios over a subset of input conditions (AR_LB*_); see main text. Links to all puzzle project pages, which include in-game scores, viewing of all submitted designs, and experimental summaries made available to the player community are compiled in **Supplemental Table 2.**

### High-throughput characterization of designs

The quantitative characterization and analysis of RNA designs through RNA-MaP was performed as previously described.^14,38,39^ DNA templates for designs were purchased in oligonucleotide pools (CustomArray, Bothell, WA), amplified by PCR or emulsion PCR, and sequenced on Illumina MiSeq instruments (primers in ref.^14^); the RNA was transcribed directly on the MiSeq sequencing chip in a repurposed Illumina Genome Analyzer II instrument. The sequences and protocols for preparing an array of clonal RNA clusters and for preparing fluorescently labeled MS2 coat protein were those described in ref.^14^. Here, several fluorescent RNA reporters were also used to measure the affinity across several input conditions. For each experiment, a full binding curve was collected for each cluster over a concentration range of 0.7–1500 nM for MS2 protein and of 0.09–1500 nM for fluorescent RNA oligos, enabling a maximum range of activation ratios of approximately 1000. For RNA input and output sequences used in the experiments, see **Supplemental Table 3**. For experimental conditions tested for Challenge 3 on RNA-MaP, see **Supplemental Table 4**.

### Independent assessment of TB-score sensors using flow cytometry

Flow cytometry enabled characterization of selected sensors across a large collection of input RNA concentrations. For each RNA sensor, DNA primer oligos for assembly were found using Primerize^62,63^ and ordered from Integrated DNA Technologies (IDT). Full-length DNA templates were assembled using standard PCR assembly protocols available at https://primerize.stanford.edu. Briefly, 100 µL of 1x PCR mix containing Phusion DNA polymerase (Thermo Fisher Scientific) was prepared with 2 µM of first and last primers (P1 and P4 or P6 for BC_AK2.2 or HP_MCTS130_DEC respectively), and 40 nM of the other primers. Then the DNA was amplified and transcribed to RNA as previously described^14^. For the MCTS_DNA_DEC construct, the sequence was ordered as a single-stranded DNA from IDT. The sequences are listed in **Supplemental Table 3**. Nucleic acid beads were prepared as in a previous study on small molecule sensors.^14^ Nucleic acids were loaded onto the magnetic bead by first preparing 3.33 𝜇L of bead mix solution by mixing 0.33 𝜇L of poly-T-coated beads, 0.175 𝜇L RNA (250 nM), and 2.825 𝜇L of H_2_O and incubated at 37°C for 5 minutes and then put on ice for 5 minute. The buffer was removed and the beads were washed three times with 100 𝜇L solution containing 1x Other buffer and 1x TMK buffer (10x Other buffer contains 1 mg/mL BSA, 10 mM DTT, 0.1 mg/mL yeast tRNA, 0.1% Tween-20; 5x TMK buffer contains 500 mM Tris-HCl pH 7.5, 400 mM KCl, 20 mM MgCl_2_). After washing, the beads were resuspended in 3.33 𝜇L of H_2_O. The bead mix is then added to 20 𝜇L TMK buffer, and 10 𝜇L Other buffer, 3 𝜇L reporter (R) RNA, and 43.66 𝜇L of H_2_O resulting in 80 𝜇L of solution. 20 𝜇L of solution containing different concentrations of RNA A, B, and C was added. The final concentration of R was 30 nM, set slightly higher than the 25 nM simulated reporter concentration in Eterna based on empirical calibration of sensor affinities from RNA-MaP experiments. Each sample was analyzed using a Sony SH800S Cell Sorter and data for 10,000 events were collected per sample. Beads were excited using a 561 nm laser and their emitted fluorescence was measured from the 600±60 nm emission channel.

### Nucleologic

Nucleologic is a Monte Carlo tree search (MCTS) algorithm^56,57^ for designing riboswitches, available at https://eternagame.org/about/software. Inspired by the Eterna player strategy of domain mapping secondary structure design (DMSSD), the sensor is treated as an ordered list of domains with each domain containing its own sequence. The root node of the MCTS is generated based on the user input. Possible inputs and outputs are limited to aptamers and RNA/DNA. When using aptamers in Nucleologic the sequence, secondary structure, and *K*_d_ must be specified. The input file must also specify the condition(s) that the sensor must satisfy in order for it to be considered a solution. Each condition is specified as ON or OFF depending on whether the output is bound or not; as well as the criterion of success, at which point the algorithm terminates even if it has not completed the total number of iterations *N*_iterations_. For example, the ON state could be input A is 100 nM and the OFF state could be input A is 0 nM; and the early termination success criterion could be specified as having an activation ratio of greater than 50. At least one ON and one OFF condition must be specified. Extra parameters to alter the MCTS run can also be specified such as the number of iterations, number of children generated, folding package (e.g., NUPACK, etc), and more. The code documentation includes details and examples of input files. The MCTS then involves growing a tree whose nodes represent sequence solutions for the sensor, with scores updated through the following four-step process:

*Step 1: Selection.* Starting from the root node of the tree, child nodes are successively chosen until a leaf node is reached. If a node has multiple children nodes, the child node that has the maximum value of the UCT (Upper Confidence bound for Trees) score is chosen.

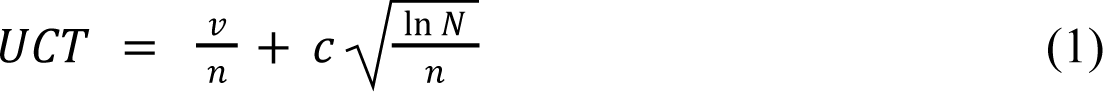

Here *N* is the total count of visits for the parent node, *n* is the total count of visits for the child node, *c* is the exploration constant, and *v* is the value of the child node’s sequence. The value *v* is defined to be sum of Boltzmann probabilities 𝑝_𝑖_ that each state matches its target condition (ON or OFF), i.e.,

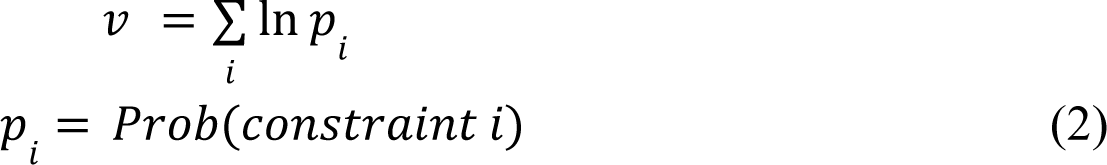

Thus the value of each node reflects the probability to fulfill the desired constraints with a higher value being close to ideal. This formulation of the node value for MCTS is heuristic and alternative formulations could be explored, but are not discussed here.

*Step 2: Expansion.* Once a leaf node is chosen, a child node is created by running the Metropolis–Hastings algorithm. Starting with the leaf node sequence, the sequence is randomly mutated through either a domain mutation or domain move (Fig. 7) and its corresponding value *v* is computed. The probability of accepting the sequence as the child node is computed as

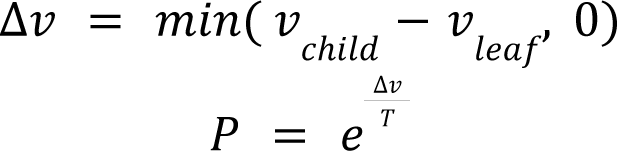

where *T* is an effective temperature. If the value of the child node candidate sequence is better (higher) than the leaf node, the sequence is always accepted. This is repeated until *n* child nodes are created from the leaf node.

*Step 3: Simulation.* From the children created, choose a random child node and compute its value using formula (2).

*Step 4: Backpropagation.* From the child chosen in step 3, backpropagate up the tree by updating the UCT score of the parent node, its parent node, etc., back up to the root node.

Typical values for MCTS searches were *n* = 3, *c* = 1*, N_iterations_* = 300 (corresponding to maximum number of expansion steps of 100), and *T* = 0.61597.

### Functions computed by RNA sensors

Equilibrium sensors can be modeled using simple equilibrium expressions involving ratios of polynomials with positive coefficients, also called positive rational polynomials, as described in the **Supplemental Appendix 2**. We give four examples from each of the challenges in this study below.

*Pilot challenge: RNA sensors for single-input oligonucleotides.* With a single input RNA A and an output reporter R, a two state model is adequate for describing the desired sensor S. As an example, the scheme for an OFF sensor with high enough concentrations of A and R is described by the equilibrium A•S ↔ R•S, and the fraction of sensor with reporter bound 𝑓 is:

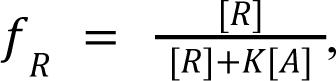

where *K* is an equilibrium constant, which is simulated in Eterna with NUPACK. At a fixed reporter concentration (100 nM simulated in Eterna), the sensor’s fluorescence switches from 1 to 0 as [A] increases from 0 to high concentrations. To precisely characterize the sensor RNA-MaP experiments, the concentration of R was titrated and its apparent dissociation constant *K^app^_d_* was measured at zero and high [A]. At equilibrium, the expression above and the standard 𝑑 relationship defining the dissociation constant, i.e., 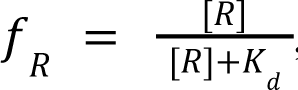, guarantees that *K^app^_d_* = *K*[A] for an accurately designed single-input sensor. Although the maximal activation ratio for the system is unbounded in this simple two-state model, taking into account other states (e.g., free sensor in the absence of A or R) leads to the maximum AR given by the ratio of the test concentration [A] (here, 100 nM) and the intrinsic hybridization affinity of A for its complement (which can be femtomolar or smaller); see ref.^48^. This value can be very large (>10^6^) so in practice, the maximum AR value for ‘binary’ sensors responding to oligonucleotide inputs is limited by the experimental range of *K^app^_d_* measurable in RNA-MaP (∼1000). Similar expressions 𝑑 and considerations hold for an ON sensor.^48^

*Challenge 1: Two-input logic gates.* For two inputs and complex logic gates, a few-state model remains sufficient to describe the sensor. The most complex case is an XOR system that responds to two inputs A and B, which is minimally described by four states: S ↔ A•R•S ↔ B•R•S ↔ A•B•S. The output of the system is described by:

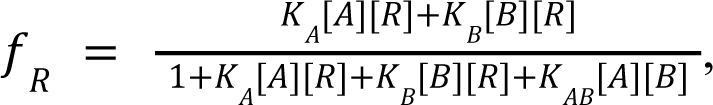

where *K_A_*, *K_B_*, and *K_AB_*are equilibrium constants that can be estimated in packages like NUPACK. For a given [R], the expression is zero under conditions without A or B, or conditions with high concentrations of both A and B, but approaches 1 with high concentrations of just A or just B. RNA-MaP experiments varied [R] to enable more precise characterization, fitting an apparent dissociation constant which, in the case of a well-designed sensor, will conform to the rational polynomial 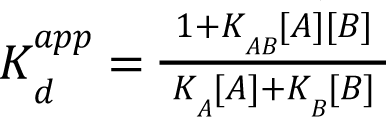. This should lead to a weak (large *K^app^_d_*) value without any inputs or with both inputs, and a tight (small *K^app^_d_*) value with high concentrations of any 𝑑 single input. For A and B that bind their complement tightly, maximal activation ratios can be >10^6^; as with single-RNA input gates, the maximum activation ratios are set here by the experimental range of 𝐾 measurable by RNA-MaP, about 1000 with MS2 protein binding. See 𝑑 **Supplemental Appendix 2** for a graph of an XOR sensor response.

*Challenge 2: Ratio sensor.* A ratio sensor requires only two states to describe, A•R•S ↔ B•S. The output of the system is:

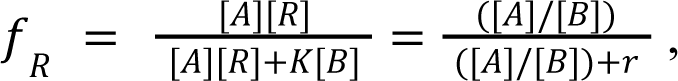

where *K* is an equilibrium constant, and *r* = *K*/[R]. At a fixed reporter concentration [R], the system therefore varies from 0 to 1 as a monotonic function of the ratio [A]/[B], with the midpoint (‘separatrix’) set by *r*, here targeted as *r* = ¼ for [R] set to the *K*_d_ for MS2 coat protein reporter; in Eterna, this condition is equivalent to simulating whether the MS2 hairpin is displayed or not displayed in the lowest free energy sensor state at the different [A] and [B]. RNA-MaP experiments characterized sensors by titrating [R] and measuring apparent dissociation constants which, for an accurately designed sensor, is given by *K^app^_d_* = *K* [A]/[B]. For a perfect sensor, a lower bound on the activation ratio is set by the two conditions whose [A]/[B] are closest to the separatrix *r* from above and below: AR_LB_ = min_[A]/[B]>*r*_ ([A]/[B]) / max_[A]/[B]<r_ ([A]/[B]).

*Challenge 3: OpenTB sensor.* Sensors computing the TB-score, which depend on concentrations of three RNA segments as [A][B]/[C]^2^, can be achieved with designs that populate just two states. For a DEC sensor, the two states are A•B•S ↔ C•C•R•S, and

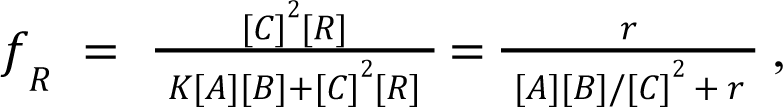

where *K* is an equilibrium constant, and *r* = [R]/*K*. Similar to the ratio sensor above, at a fixed reporter concentration [R], the system varies from 1 to 0 as a monotonic function of the ratio [A][B]/[C]^2^ with the midpoint (‘separatrix’) set by *r*. Here, we targeted *r* = 1/16 at [R] = 25 nM. This value for *r* was set based on a study^34^ that defined the TB-score as (GBP5 + DUSP3) / 2 − KLF2, with individual values defined as logarithm (base 2) of gene concentrations to enable convenient comparison to quantitative RT-PCR cycle threshold values, and a separatrix value of −2 based on clinical samples. Again, similar to the ratio sensors, RNA-MaP experiments read out *K^app^_d_* = *K*[A][B]/[C]^2^ for a perfect TB-score sensor, and AR_LB_ = min_[A][B]/[C][C]>*r*_ ([A][B]/[C]^2^) / max_[A][B]/[C][C]<r_ ([A][B]/[C]^2^). Similar expressions and considerations hold for an INC sensor (reporter binding at [A][B]/[C] above rather than below a threshold).

## DATA AVAILABILITY

Experimental data for all figures, including estimated *K*_d_ values for each tested sequence, can be found in the Github repository: https://github.com/eternagame/paper-data-rationally-designed-RNA-sensor. Data compiled at the finer level of individual RNA-MaP sequence clusters are available at https://github.com/eternagame/EternaDataRibonet and described in reference.^64^

## CODE AVAILABILITY

Code for generating simulated plots for visualizing response curves to different concentrations of A, B, and C can be found in the Github repository: https://github.com/eternagame/conc_plots. Code for Nucleologic can be downloaded from https://eternagame.org/about/software.

## AUTHOR CONTRIBUTIONS

R.D., P.K., and W.J.G. conceptualized the project. F.P. and J.J.N. wrote code for, designed, and set up puzzles on Eterna. R.W.-O. coordinated research on Eterna. J.O.L.A., R.D., and W.J.G. built RNA-MaP instruments. Eterna Participants created designs and analyzed data. J.O.L.A. and W.K. performed RNA-MaP experiments. M.J.W. and J.O.L.A. analyzed Eterna designs and RNA-MaP data. F.M., C.A.C., W.K., and M.G. designed and performed the flow cytometry experiments. C.A.C. created Nucleologic. M.J.W., J.O.L.A., F.M., H.K.W.-S., and C.A.C. prepared figures. C.A.C., F.M., H.K.W.-S., R.D., and W.J.G. wrote the manuscript, with all authors providing comments.

## ETHICS DECLARATION

The authors declare no competing interests.

## ACKNOWLEDGEMENTS

At Stanford, we thank F. Sun for technical assistance; C. Layton for expert advice in building the MaP instrument; M. Gotrik for advice on flow cytometry experiments; and T. Sweeney for discussion of the TB-score. J. Anderson-Lee, Boris Rudolphs and E. Fisker for discussion of Eterna designs. This work was supported by a Stanford Graduate Fellowship (to C.A.C.), Stanford School of Medicine Discovery Innovation Award (to R.D.), a Burroughs-Wellcome Foundation Career Award (to R.D.), JIMB Seed Grant (to R.D. and W.J.G.), NIH Grants R01 GM100953 (to R.D.), R35 GM122579 (to R.D.), the Howard Hughes Medical Institute (HHMI, to R.D.), R01 GM111990 and P50 HG007735 (to W.J.G.), and Bill and Melinda Gates Foundation (GH-VAP ID-28 to R.D. and P.K; OPP1113682 to P.K.). M.J.W. was supported by NSF Graduate Research Fellowship DGE-114747, NLM Biomedical Informatics Training Grant T15 LM007033, and NIH Ruth L. Kirschstein National Research Service Award F31GM125151.

W.J.G. acknowledges support as a Chan-Zuckerberg Investigator. Computational design was performed on the Stanford BioX^3^ cluster, supported by NIH Shared Instrumentation Grant S10 RR02664701. This article is subject to HHMI’s Open Access to Publications policy. HHMI lab heads have previously granted a nonexclusive CC BY 4.0 license to the public and a sublicensable license to HHMI in their research articles. Pursuant to those licenses, the author-accepted manuscript of this article can be made freely available under a CC BY 4.0 license immediately upon publication.

