## Extended Figures for "Compact RNA sensors for increasingly complex functions of multiple inputs"

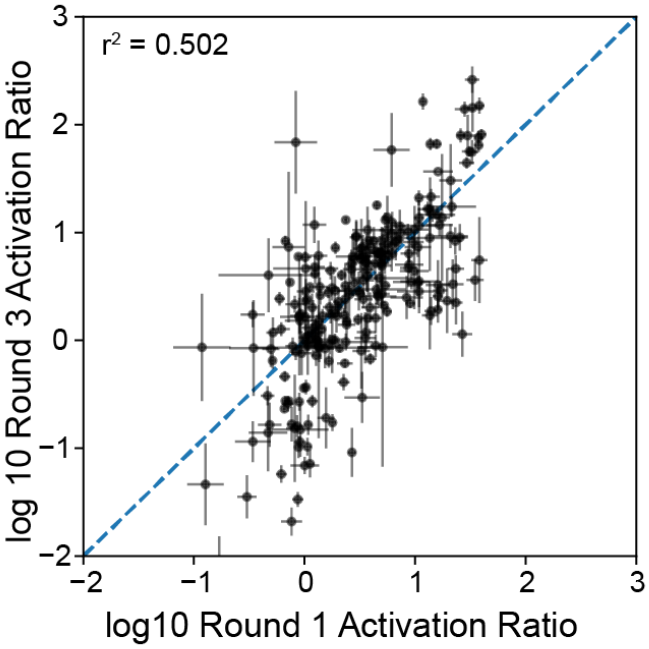


**Extended Data Figure 1. Reproducibility of activation ratio estimation with RNA-MaP experiments.** In the Pilot Challenge (single-input RNA sensors), 245 designs for Template 1 Round 1 were experimentally re-evaluated in Round 3. Designs shown were for Template 1 (Figure 2) involving RNA-input and MS2-output. Pearson correlation coefficient is $r^2 = 0.502$.

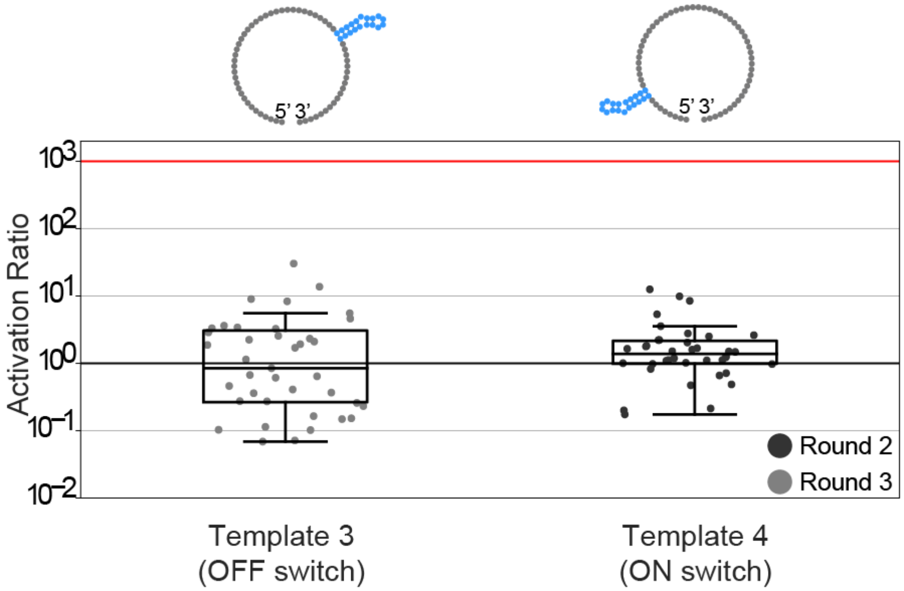


**Extended Data Figure 2. Puzzle results for Templates 3 and 4 in later rounds of single-input RNA sensors.** Template 3 submissions were from Eterna players in their third round of this challenge. Template 4 submissions were from an invitation-only Eterna puzzle for Stanford students of the graduate course Biochemistry/Biophysics/Structural Biology 241 (Round 2 supplemental in Supplemental Table 2). The activation ratios in these separate later rounds did not exceed activation ratios achieved by Eterna participants in early rounds. The red horizontal line is the approximate maximum activation ratio that can be experimentally measured in RNA-MaP (1000).

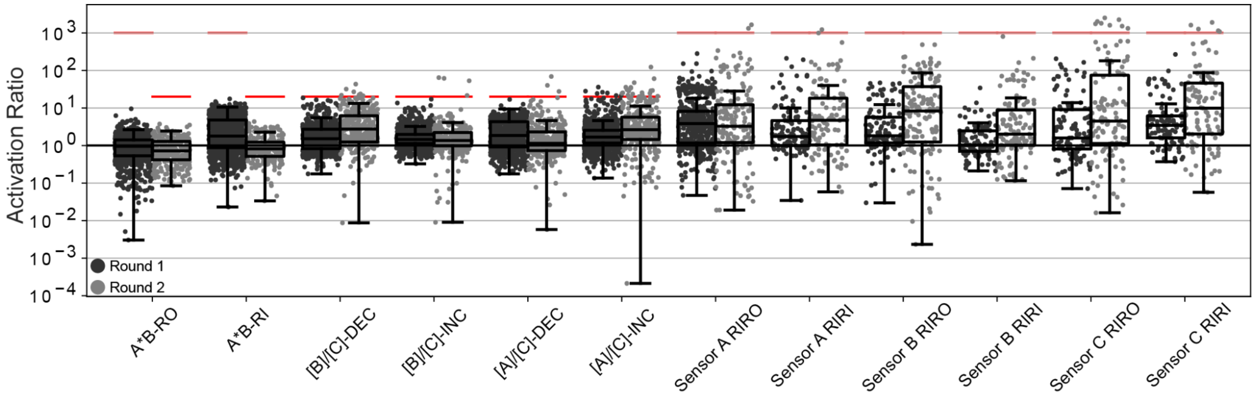


**Extended Data Figure 3. Additional puzzles presented alongside Challenge 3 (OpenTB sensor).** In addition to the two puzzles that correspond to designing a DEC or INC sensor for the three-gene TB-score, 12 additional puzzles were presented to players as sub-problems.The rounds correspond to the same rounds as in Figure 5 (see Supplemental Table 2 for sequences of A, B, C, and reporter R). Activation ratios (lower bound) are computed using a subset of the ON and OFF conditions related to actual conditions simulated in the Eterna puzzles (denoted as $AR_{LB*}$ in the main text). For A*B puzzles, in Round 1, four conditions were used for scoring (0 nM A&B, 0 nM A & 100 nM B, and 100 nM A & 0 nM B; and 100 nM A & B) and, in Round 2, three conditions were used for scoring (5 nM A&B, 100 nM A & 5 nM B; and 100 nM A & B). For the ratio sensor puzzles, two conditions were used for scoring with 5 vs. 100 nM of the 'numerator' oligonucleotide input, and 100 nM C in both conditions. For single-input sensor puzzles, two conditions were used for scoring, 0 nM or 100 nM for the input, and 0 nM concentrations of other oligonucleotides. See also Supplemental Table 5. The red horizontal lines are the approximate maximum activation ratio that can be experimentally measured in RNA-MaP (1000); when the red line is below 1000 it indicates the theoretical maximum $AR_{LB*}$ based on the conditions for the respective puzzles. Abbreviations: INC or RI, puzzles where binding of output oligonucleotide R occurs at high values of input concentrations; DEC or RO, puzzles where binding of output occurs at low input values.

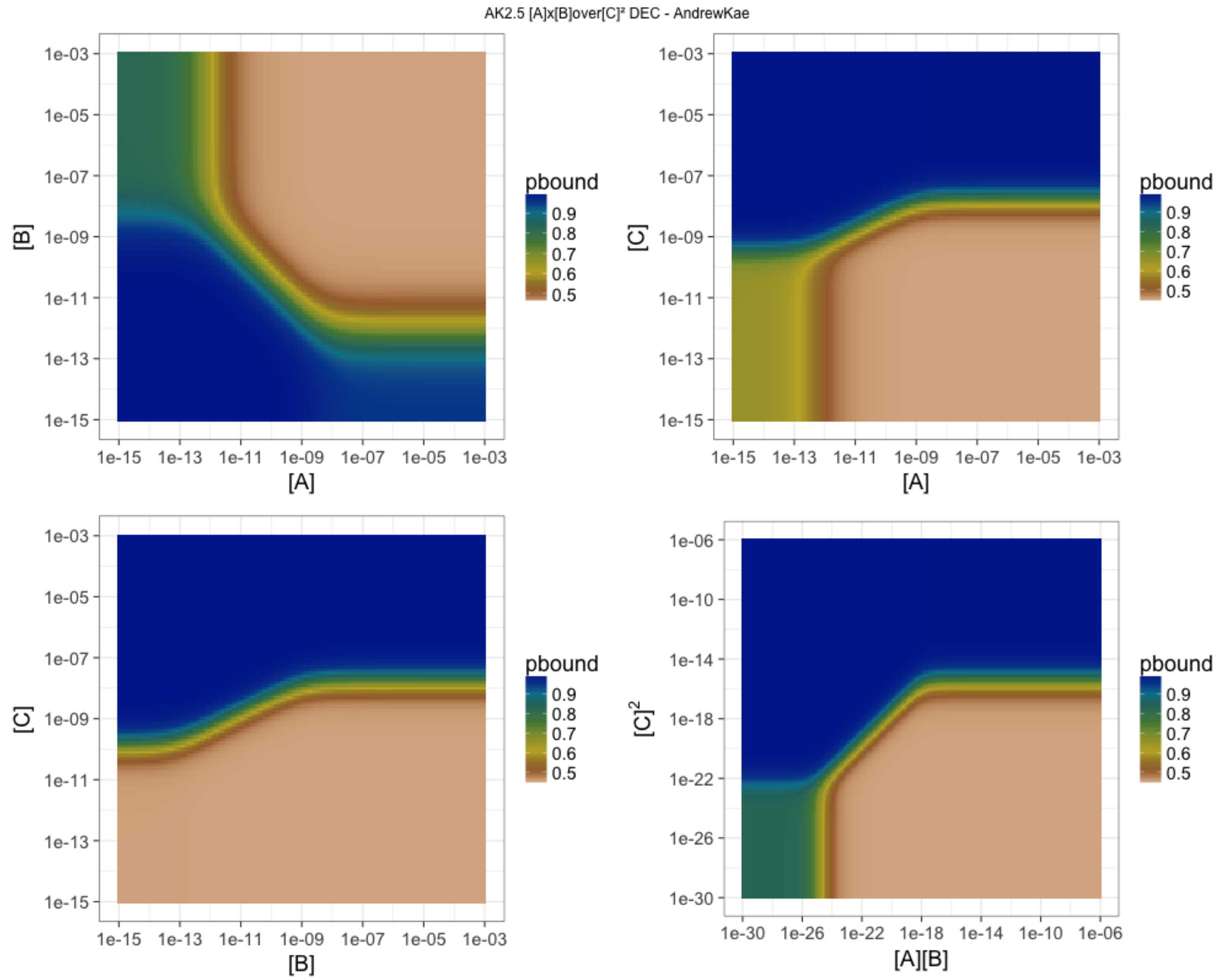


**Extended Data Figure 4. On-line simulation plots for design AK2.5 submitted by AndrewKae.** Simulation plots, made available to Eterna participants through a separate webserver (code available at https://github.com/eternagame/conc_plots), showed the probability of binding the output reporter RNA at various concentrations of A, B, and C (given in M) as predicted by NUPACK. The simulated reporter RNA concentration was 1 nM. [C] = 1 nM in [A] vs [B]. [B] = 1 nM in [A] vs [C]. [A] = 1 nM in [B] vs [C].

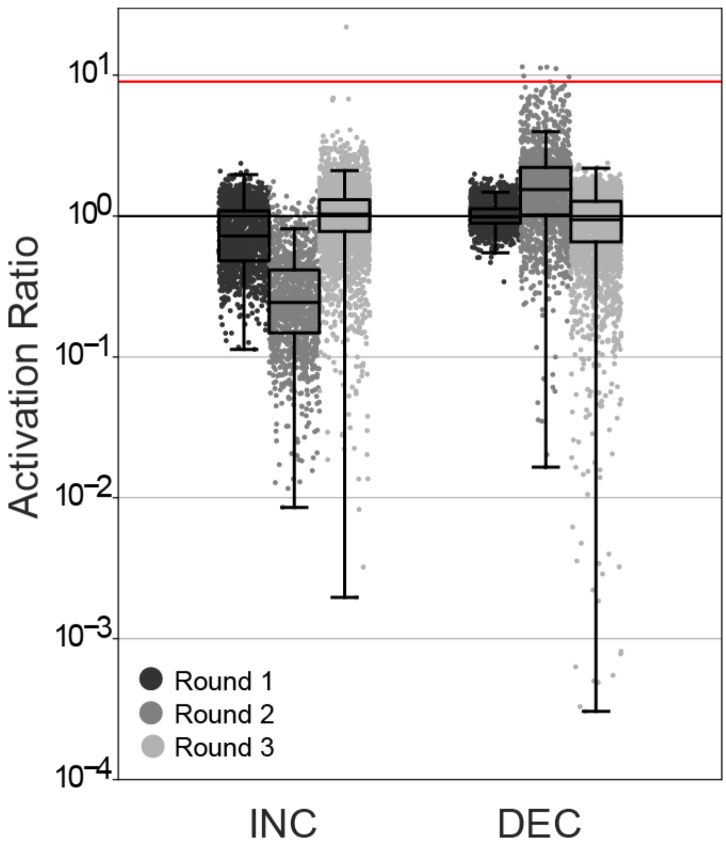


**Extended Data Figure 5. Activation ratios computed over four special conditions for Rounds 1 through 3 of Challenge 3: OpenTB sensors.** Activation ratio shown is the lower bound with respect to the 4 conditions presented to players in Eterna, rather than over all conditions tested (12 or more; see Supplemental Table 5). These $AR_{LB*}$ values also informed scores returned to Eterna participants as feedback during the OpenTB challenge. The four conditions are (1) 100 nM A & 100 nM C, (2) 100 nM B & 100 nM C, (3) 50 nM A&B & 300 nM C, (4) 50 nM A&B & 100 nM C. The red horizontal line is the theoretical maximum ($AR_{LB*} = 9$) for these four conditions.

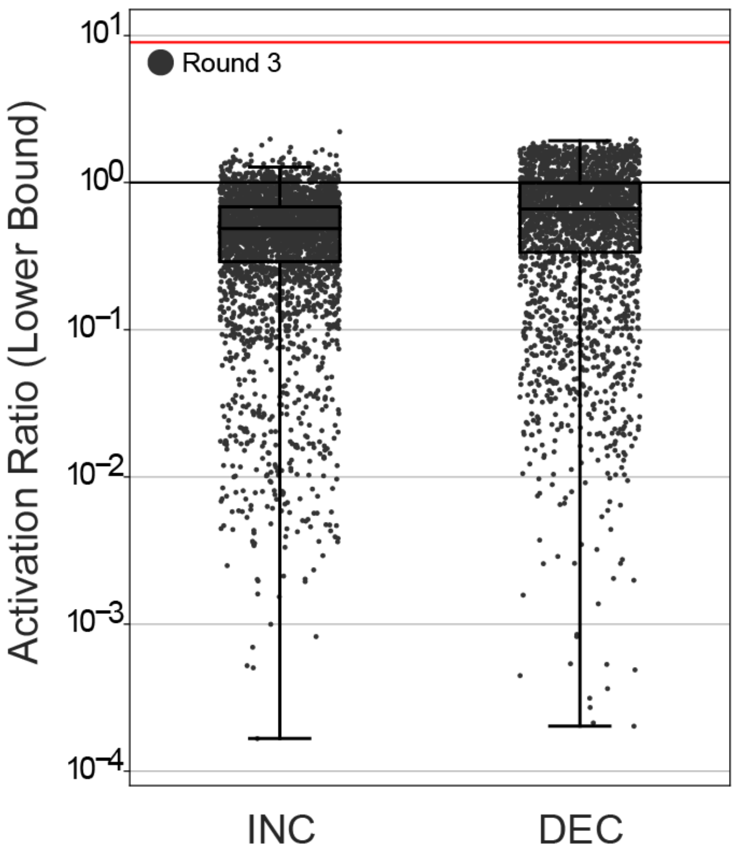


**Extended Data Figure 6. Activation ratio (lower bound) Round 3 of Challenge 3: OpenTB sensors**. Round 3 reverted input and output RNA oligonucleotide sequences back to Round 1 sequences, and performed similarly poorly to Round 1, compared to Round 2. Activation ratio shown is the lower bound over all measured conditions. The red horizontal line is the theoretical maximum ($AR_{LB}$ = 9).

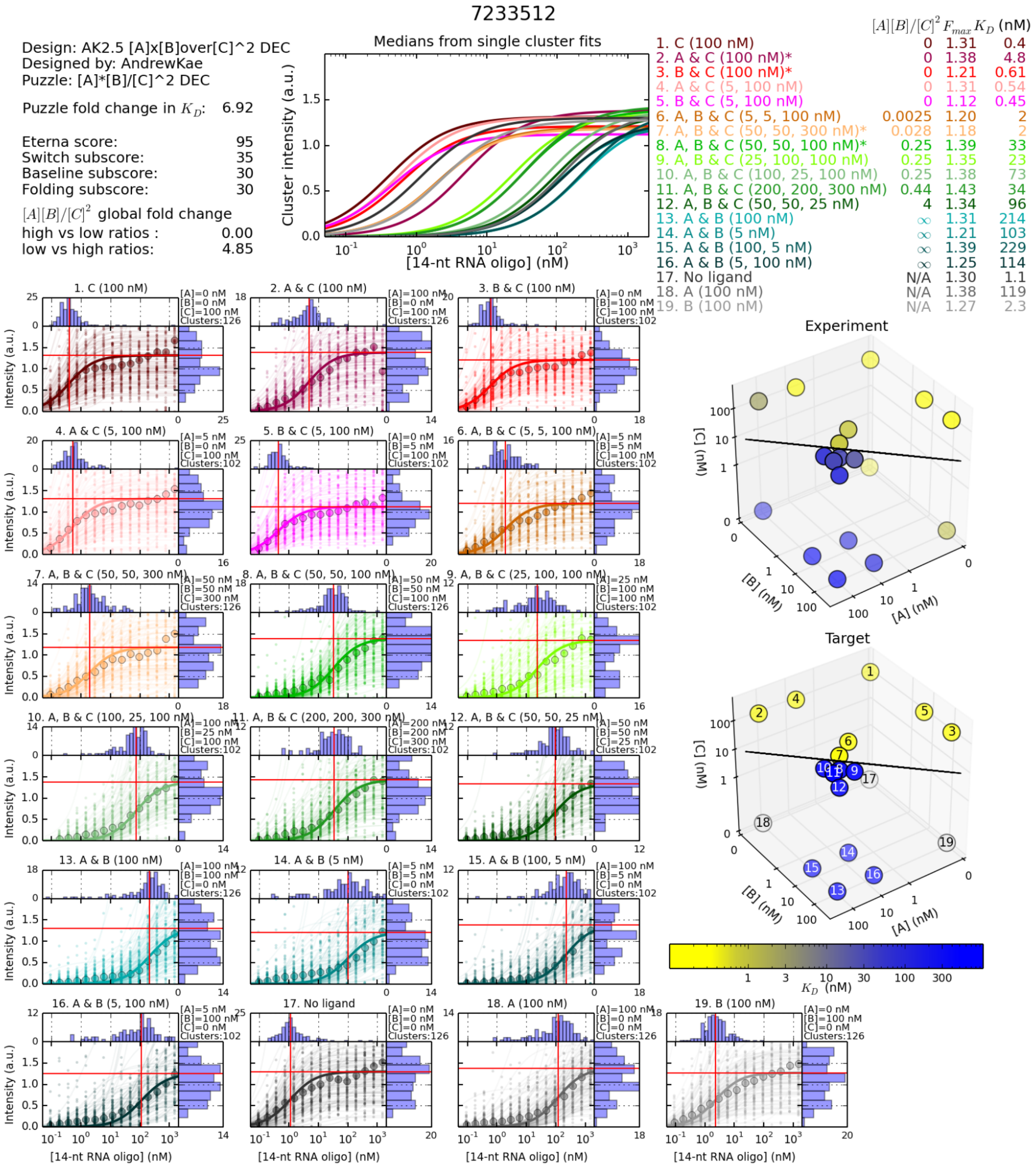


**Extended Data Figure 7. Example of RNA-MaP data summary plot made available to Eterna participants for OpenTB challenge.** These data are for Round 2 of Challenge 3: OpenTB sensor, for design AK2.5 submitted by AndrewKae. The affinity of the designed RNA to the reporter RNA is computed as various conditions.

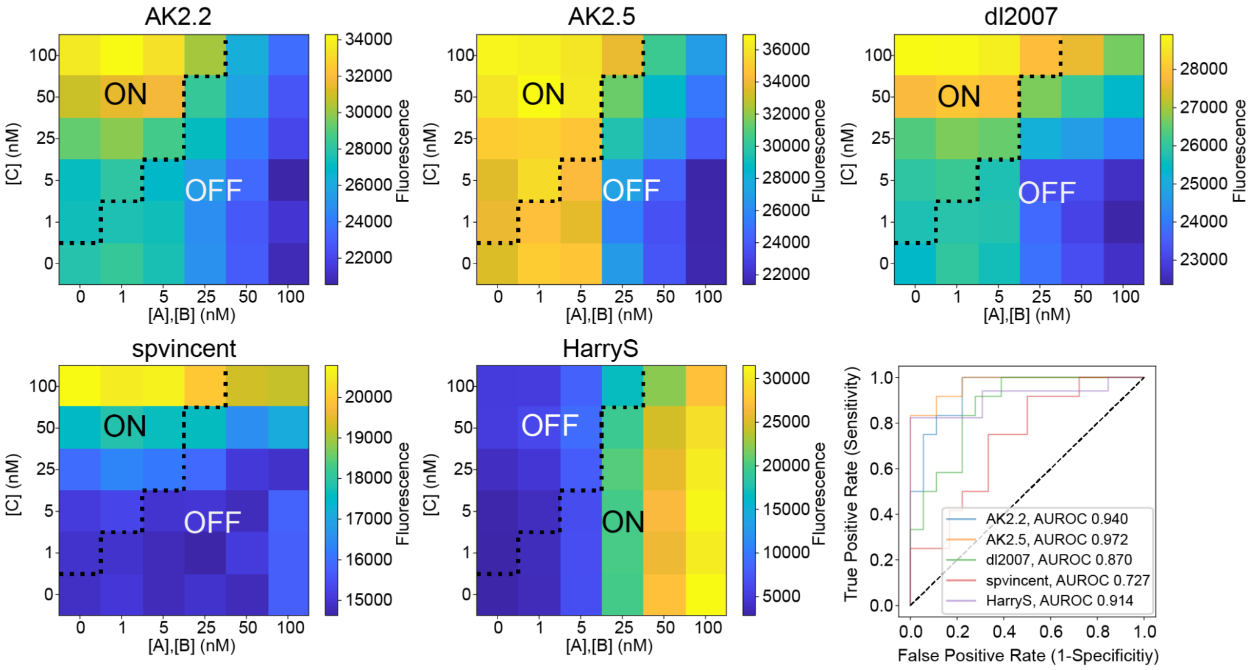


**Extended Data Figure 8. Flow cytometry characterization of top player designs from Round 2 of Challenge 3: OpenTB calculators.** Five different player designs were tested in 36 different conditions as shown in the heat map. The title of the heatmap are the player names except AK2.2 and AK2.5 which are designs from Eterna player AndrewKae. From left to right and top to bottom the 5 designs are from (1) player: AndrewKae design ID: 7233501 design name: "AK2.2 [A]x[B]over[C]^2 DEC", (2) player: AndrewKae design ID: 7233512 design name: "AK2.5 [A]x[B]over[C]^2 DEC", (3) player: dl2007 design ID: 7172797 design name: "A*B/C^2 DEC 8  9", player: spvincent design ID: 7230841 design name: "A*B/C*C Dec Mod20", player: HarryS design ID: 7252060 design name: "[A]*[B]/([C]*[C]) - INC  Attempt #68 (Inspired by Eli Fisker)". Designs 1-4 are DEC sensors and design 5 is an INC sensor. Receiver operating characteristic (ROC) curves of all five designs are shown on the bottom right.

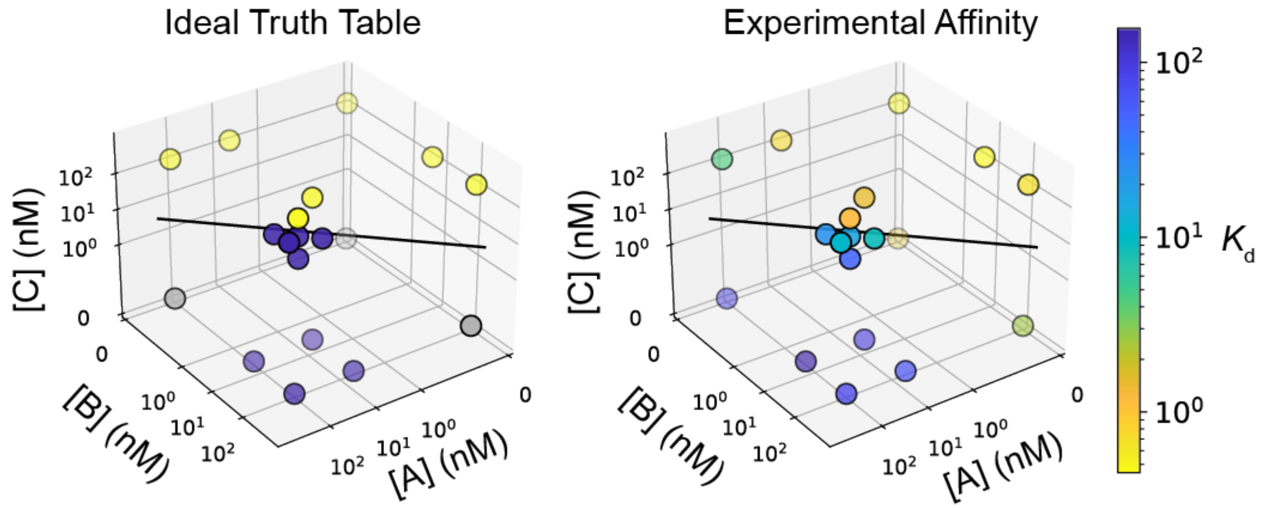


**Extended Data Figure 9. 3D truth table for design AK2.2 from Round 2 of challenge 3: OpenTB calculator.** 3D scatter plot of the ideal truth table (left) and experimental values (right). The separatrix plane for $[A][B]/[C]^2 = 1/16$ appears as a line due to the chosen view of the 3D plot, and gray points in ideal truth tables are undefined with respect to the three-gene TB-score.

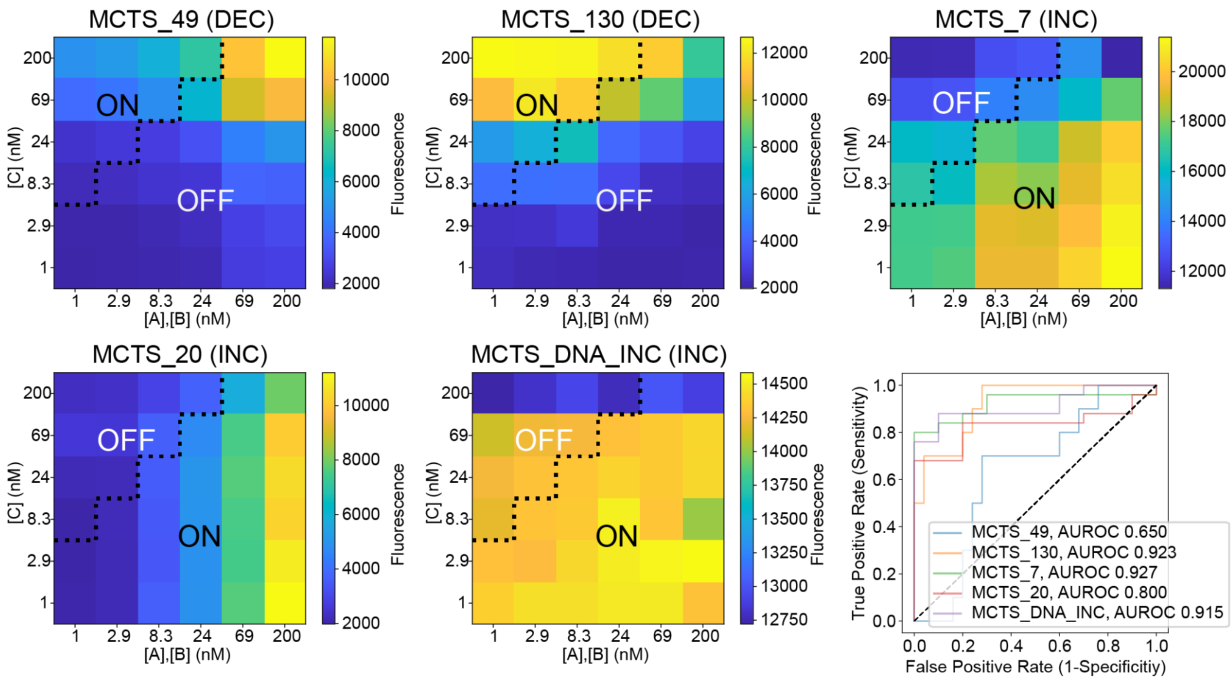


**Extended Data Figure 10. Flow cytometry characterization of other Nucleologic designs.** The top two DEC and INC RNA designs as well as an INC DNA design from Nuclelogic were tested in 36 different conditions as shown in the heat map. The title of the heatmap is the design name and the type of sensor INC or DEC is in parentheses. Receiver operating characteristic (ROC) curves of all designs are shown on the bottom right.
