## Supplemental Appendix 2 for "Compact RNA sensors for increasingly complex functions of multiple inputs"

### Supplementary Appendix 2: Molecular rational function approximation

#### 1 Single molecules as rational functions

Nature exhibits remarkable computational abilities mediated through the complex interactions between nucleic acids and proteins. The capacity of a molecule to bind input and output ligand(s) endows the potential to carry out computations in-situ. Among the vast array of biomolecules, nucleic acids, such as DNA and RNA, hold particular promise as molecular calculators due to their versatility and the ease with which they can be designed and manipulated. Although the complexity of molecular calculators can be enhanced by linking multiple calculators together, this supplement concentrates on the investigation of single-molecule calculators.

##### 1.1 Partition function of a generalized molecular calculator at equilibrium

Given a single-molecule calculator, $S$, that has $N$ unique binding sites for ligand $A$ and $M$ unique binding sites for a reporter ligand $R$ we can write out all the states of S complexed with $A$ and $R$. We denote $S{\cdot}A_k$ as the set of all states where $A$ bound to the $k$th-unique binding site on S. We let $R$ denote a fluorescent reporter that gives a signal of 1 when bound by $S$ and a signal of 0 when free in solution. In practice, other output modalities can be used such as transcription activation.

$$\text{Let } S_A = \{A_i | i = 1..N\}, S_R = \{R_i | i = 1..M\}$$

$$\binom{S_i}{k} \text{ denote the set of all combinations choosing } k \text{ from } S_i$$

$$\text{E.g.} \binom{S_A}{2} = \bigcup_{i=1}^{N} \bigcup_{j=i}^{M} A_i{\cdot}A_j = \{A_i{\cdot}A_j | i = 1..N, j = i..M\}$$

We can express all the states of $S$ complexed with $A$ and $R$.

$$\begin{aligned}
\text{States } &= \{S \text{ bound to 1 copy of } A\} \cup \{S \text{ bound to 2 copy of } A\} \cup ... \\
&= \left\{ \bigcup_{j=0}^{M} \bigcup_{a \in \binom{S_A}{0}} \bigcup_{r \in \binom{S_R}{j}} S{\cdot}a{\cdot}r \right\} \cup \left\{ \bigcup_{j=0}^{M} \bigcup_{a \in \binom{S_A}{1}} \bigcup_{r \in \binom{S_R}{j}} S{\cdot}a{\cdot}r \right\} \cup ... \\
&= \bigcup_{i=0}^{N} \bigcup_{j=0}^{M} \bigcup_{a \in \binom{S_A}{i}} \bigcup_{r \in \binom{S_R}{j}} S{\cdot}a{\cdot}r \qquad (1)
\end{aligned}$$

From the set of states, we can proceed to computing the partition function $Z$. Assume the system is in thermodynamic equilibrium and $[R]$ is constant, we may write the partition function of the system as

$$\begin{aligned} Z &= \sum_{i=0}^{N}\sum_{j=0}^{M}\sum_{a\in\binom{S_A}{i}}\sum_{r\in\binom{S_R}{j}} Z_{S\cdot a\cdot r} \\ &= \sum_{i=0}^{N}\sum_{j=0}^{M}\frac{[A]^i[R]^j}{K_A^iK_R^j}e^{-\beta\Delta G_{i,j}} \\ &= \sum_{i=0}^{N}[A]^i\sum_{j=0}^{M}\frac{[R]^j}{K_A^iK_R^j}e^{-\beta\Delta G_{i,j}} \end{aligned}$$

where $\beta = 1/k_BT$, where $k_B$ is Boltzmann's constant and T is temperature. $K_A$ and $K_R$ are the dissociation constants of $A$ and $R$ respectively with units $\frac{L}{mol}$

Let $b_{i,j} = \frac{[R]^j}{k_{i,j}}e^{-\beta\Delta G_{i,j}}$

$$Z = \sum_{i=0}^{N}[A]^i\sum_{j=0}^{M}b_{i,j} \tag{2}$$

The term $b_{i,j}$ represents the degeneracy and energetic contributions of all states of $S$ complexed with $i$ copies of $A$ and $j$ copies of $R$.

We can simplify (2) further by letting $x = [A]$ and $c_i = \sum_{j=0}^{M} b_{i,j}$ resulting in $Z = \sum_{i=0}^{N} c_ix^i$

#### 1.2 Calculating the output probability of the biomolecule

Using the partition function defined in the previous section, we can compute the probability of n copies of $R$ bound to the $S$ which is related to the observable in an experiment.

We write for our partition function of activated states,

$$Z_{\text{out}} = \sum_{i=1}^{N} b_ix^i \tag{3}$$

Since the set of activated states is always a subset of the total states represented by $c_i$ we have $0 \leq b_i \leq c_i$. Thus the signal, $f(x)$ of the system is represented by $Z_{\text{out}}/Z$.

If $S$ can only bind up to one copy of the reporter we can write out the functional equation of a system where $M = 1$. We assume the signal of the reporter is 0 when free in solution, and 1 when bound.

$$f(x) = p(R\text{ bound}) = \frac{\sum_{i=0}^{N} b_ix^i}{\sum_{i=0}^{N} c_ix^i} \leq 1,\ 0 \leq b_i \leq c_i \tag{4}$$

If $M > 1$ our function is

$$
\begin{aligned}
f(x) &= \sum_{j=1}^{M} j \cdot \frac{\sum_{i=0}^{N} b_{i,j} x^i}{\sum_{i=0}^{N} c_i x^i} \\
&= \frac{\sum_{i=0}^{N} x^i \sum_{j=1}^{M} j \cdot b_{i,j}}{\sum_{i=0}^{N} c_i x^i} \\
d_i &= \sum_{j=1}^{M} j \cdot b_{i,j} \\
f(x) &= \frac{\sum_{i=0}^{N} d_i x^i}{\sum_{i=0}^{N} c_i x^i} \leq M, \qquad (5)
\end{aligned}
$$

$$
\begin{aligned}
c_i, d_i &\geq 0 \\
d_i = 0 &\iff c_i = 0
\end{aligned}
$$

One way to interpret the expression (5) is through probability. $f(x)$ can be written as the sum of probabilities of different number of copies bound attenuated by some signaling strength factor.

$$
f(x) = \sum_{k=0}^{N} A_k p(k) \qquad (6)
$$

$A_k$ represents the signaling strength when $k$ copies are bound and $p(k) = \frac{c_k x^k}{\sum_{i=0}^{N} c_i x^i}$ is the probability of $k$ copies bound. Depending on $M$, $A_k \cdot c_k = b_k$ or $d_k$.

In theory, we can express almost any positive rational function with positive coefficients as the partition function of a molecular calculator at equilibrium (5).

One limitation is that since $d_i = 0$ if and only if $c_i = 0$ the degree of the numerator must be less than or equal to the degree of the denominator. In order to overcome this limitation one can use $\frac{1}{f(x)}$ as the output function.

#### 1.3 Molecular functions in higher dimensions

We can expand our function (7) to higher dimensional rational functions. Let the molecule, $S$, having binding sites for ligand $x_1$, $x_2$, ..., $x_n$ where each ligand $x_k$ has $N_k$ unique binding sites. Also, let $S$ able to bind $m$ copies of a reporter. We can write out the new partition function.

$$
Z = \sum_{\alpha_1=0}^{N_1} \sum_{\alpha_2=0}^{N_2} ... \sum_{\alpha_n=0}^{N_n} c_{\alpha_1,\alpha_2,...,\alpha_n} x_1^{\alpha_1} x_2^{\alpha_2} ... x_n^{\alpha_n} \qquad (7)
$$

Using this partition function we can write the molecular calculator's output signal as a higher dimensional rational function which has the same constraints as (5).

$$
f(x_1, x_2, ..., x_n) = \frac{\sum_{\alpha_1=0}^{N_1} \sum_{\alpha_2=0}^{N_2} ... \sum_{\alpha_n=0}^{N_n} d_{\alpha_1,\alpha_2,...,\alpha_n} x_1^{\alpha_1} x_2^{\alpha_2} ... x_n^{\alpha_n}}{\sum_{\alpha_1=0}^{N_1} \sum_{\alpha_2=0}^{N_2} ... \sum_{\alpha_n=0}^{N_n} c_{\alpha_1,\alpha_2,...,\alpha_n} x_1^{\alpha_1} x_2^{\alpha_2} ... x_n^{\alpha_n}} \qquad (8)
$$

### 2 Polynomial approximation

#### 2.1 Polynomial decomposition

In addition to approximating rational functions, it is also possible for the system to approximate any positive polynomial. The Stone–Weierstrass theorem [3] states that any continuous function defined on a closed interval [a, b] can be uniformly approximated as closely as desired by a polynomial function. It is possible to create a polynomial to any desired function.

Let $f(x) =$ be our polynomial where $f(x) > 0$ when $x \geq 0$. We apply these constraints on the range and domain since partition functions can only express positive values and the input in the system can never be negative. Poincaré showed that polynomials can be decomposed into linear and quadratic terms.

$$f(x) = F_1 \cdot F_2 \cdot F_3 \cdot F_4 \tag{9}$$

$$\begin{aligned} F_1 &= \prod_i x + x_{n,i} \\ F_2 &= \prod_i x^2 - a_i x + b_i \\ F_3 &= \prod_i x^2 + \alpha_i x + \beta_i \\ F_4 &= \prod_i x - x_{p,i} \\ a_i, b_i &> 0 \\ \alpha_i, \beta_i &> 0 \\ x_{n,i}, x_{p,i} &> 0 \end{aligned}$$

Since we have a positive function for $x \geq 0$, $f(x) = \cdot F_1 \cdot F_2 \cdot F_3$. Since we know all the coefficients in $F_1$ and $F_3$ are positive if we can substitute the quadratic terms in $F_2$ with rational polynomials with positive coefficients, then it is possible to express any $f(x)$ as a rational polynomial with positive coefficients.

#### 2.2 Quadratic expressed as positive rational function

Meissner [1] (include translation), simplified by Motzkin and Straus [2], showed that $F_2$ can be rewritten as a rational function with positive coefficients. Here, we briefly summarize these results.

Start with a quadratic term $x^2 - ax + b$.

$$x^2 - ax + b = \frac{Q_2(x)}{Q_1(x)} \tag{10}$$

$$\begin{aligned} Q_2(x) &= \rho^2 a_0 \left(1 - \frac{\sin(m+2)\theta}{\sin\theta} \cdot \frac{x^{m+1}}{\rho^{m+1}} + \frac{\sin(m+1)\theta}{\sin\theta} \cdot \frac{x^{m+2}}{\rho^{m+2}}\right) \\ Q_1(x) &= \sum_{k=0}^{m} \frac{a_{k+1}}{\rho^k} x^k \end{aligned}$$

$$\rho^2 = b,\ \ a = 2\rho cos\theta$$
$$m+1 < \frac{\pi}{\theta} < \frac{\pi}{\sin\theta}$$
$$a_k = \frac{a_0}{\sin\theta}\sin(k+1)\theta,\ \ k = 0,1,...,m$$
$$a_{-1} = a_{m+1} = 0$$
$$b_k = a_{k-1} + a_{k+1} - 2a_k\cos\theta,\ \ k = 0,1,...,m$$

Thus, any positive polynomial can be represented as a rational function with positive coefficients. In theory we can approximate any such polynomial using biomolecular calculators.

For example, using (10) we can transform $x^2 - 2x + 8$.

$$x^2 - 2x + 8 = \frac{\frac{1}{4}x^3 + \frac{1}{2}x^2 + 8}{\frac{1}{4}x + 1} \tag{11}$$

This is also true in the case of higher dimension polynomials which was proved by Meissner [1]. Motzkin and Straus showed that there is an extra constraint [2]. The highest degree homogenous part of the polynomial must not have any nonnegative zeros. For example, $1 + (x-y)^2$ cannot be written as a rational function with positive coefficients since $(x-y)^2$ contains nonnegative zeros.

### 3 Examples functions

#### 3.1 Sigmoid function

Let the molecule, $S$, able to bind only one copy of $A$ and one copy of $R$.

$$f(x) = \frac{b_0 + b_1x}{c_0 + c_1x} \tag{12}$$

This represents the simplest space of rational functions for the system. In addition to containing the constant functions where the output is fixed independent of the input, ON and OFF-switches are also in this space.

An example ON-switch function would be $f(x) = \frac{x}{1+x}$. If we substitute $x$ for $e^u$ we can transform our ON-switch function into a sigmoid where the domain is $(-\infty, \infty)$.

$$f(u) = \frac{e^u}{1+e^u} = \frac{1}{1+e^{-u}}$$

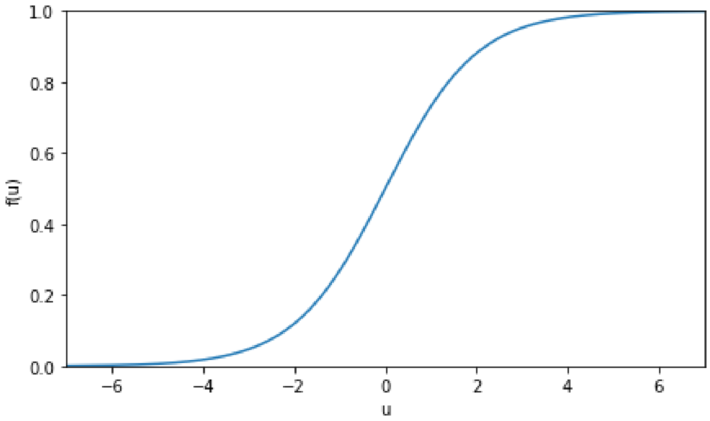


Figure 1: Example sigmoid function

If one defines an output threshold, such that responses above the threshold are labeled as “1”, and outputs below the threshold are labeled as “0”, then the sigmoid function above has two “islands” in its domain. One island corresponds to an output of 0 and the other island corresponds to an output of 1 (terminology from [4]). The number of islands can be viewed as a measure of complexity: a computer with more islands in its domain has the potential to perform more complex computations.

#### 3.2 Multi-island function

By incorporating more binding sites to the input $A$, we can create more complex functions that have multiple islands. Let the molecule, $S$, able to bind 2 copies of $A$. This allows the function to have up to three islands in its domain. Such a setup could be useful for detection of a ligand within a specific range of concentrations.

From this system we can create the following function:

$$f(x) = \frac{x}{1 + x + x^2} \tag{13}$$

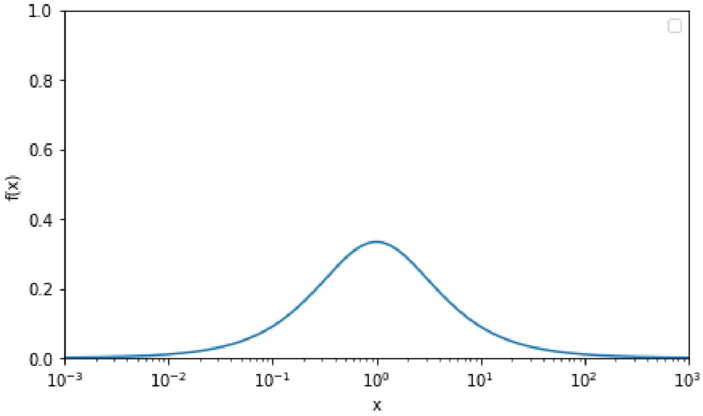


Figure 2: Example three island function

We can create more complex island functions by increasing the number of binding sites beyond 2. If the molecule, $S$, able to bind 4 copies of $A$ then five islands becomes possible.

$$f(x) = \frac{x + x^3}{.01 + x + 10x^2 + x^3 + .01x^4} \tag{14}$$

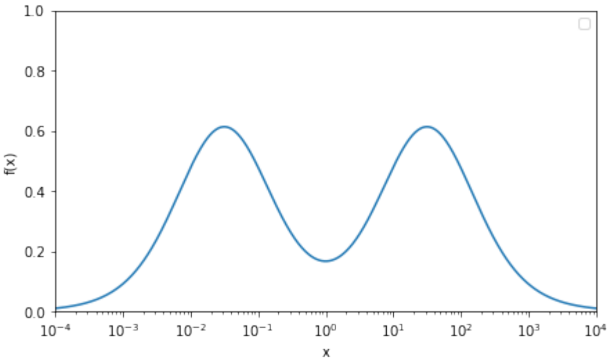


Figure 3: Example five island function

The maximum number of islands is $N + 1$ where $N$ is the number of unique binding sites for ligand $A$.

##### 3.3 Two-input logic gate

Various two-input logic gates can also be modeled by allowing for two inputs. Let $x$ and $y$ represent the concentration of two different input ligands. We can create a function for a XOR logic gate.

$$f(x, y) = \frac{1 + xy}{1 + x + y + xy} \tag{15}$$

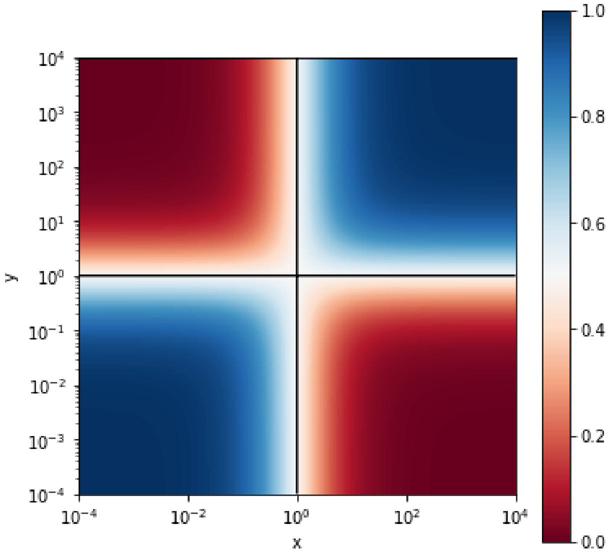


Figure 4: Example XOR logic gate function
